# Complement activation at injury sites drives the phagocytosis of necrotic cell debris and resolution of liver injury

**DOI:** 10.1101/2024.08.23.609344

**Authors:** Sofie Vandendriessche, Matheus Silvério Mattos, Emilia Laura Bialek, Sara Schuermans, Paul Proost, Pedro Elias Marques

## Abstract

Cells die by necrosis due to excessive chemical or thermal stress, leading to plasma membrane rupture, release of intracellular components and severe inflammation. The clearance of necrotic cell debris is crucial for tissue recovery and injury resolution, however, the underlying mechanisms are still poorly understood, especially in vivo. This study examined the role of complement proteins in promoting clearance of necrotic cell debris by leukocytes and their influence on liver regeneration. We found that independently of the type of necrotic liver injury, either paracetamol (APAP) overdose or thermal injury, complement proteins C1q and (i)C3b were deposited specifically on necrotic lesions via the activation of the classical pathway. Importantly, C3 deficiency led to a significant accumulation of necrotic debris and impairment of liver recovery in mice, which was attributed to decreased phagocytosis of debris by recruited neutrophils in vivo. Monocytes and macrophages also took part in debris clearance, although the necessity of C3 and CD11b was dependent on the specific type of necrotic liver injury. Using human neutrophils, we showed that depletion of C1q or C3 caused a reduction in the volume of necrotic debris that is phagocytosed, indicating that complement promotes effective debris uptake by neutrophils in mice and humans. In summary, complement activation at injury sites is a pivotal event for necrotic debris clearance by phagocytes and determinant for efficient recovery from tissue injury.

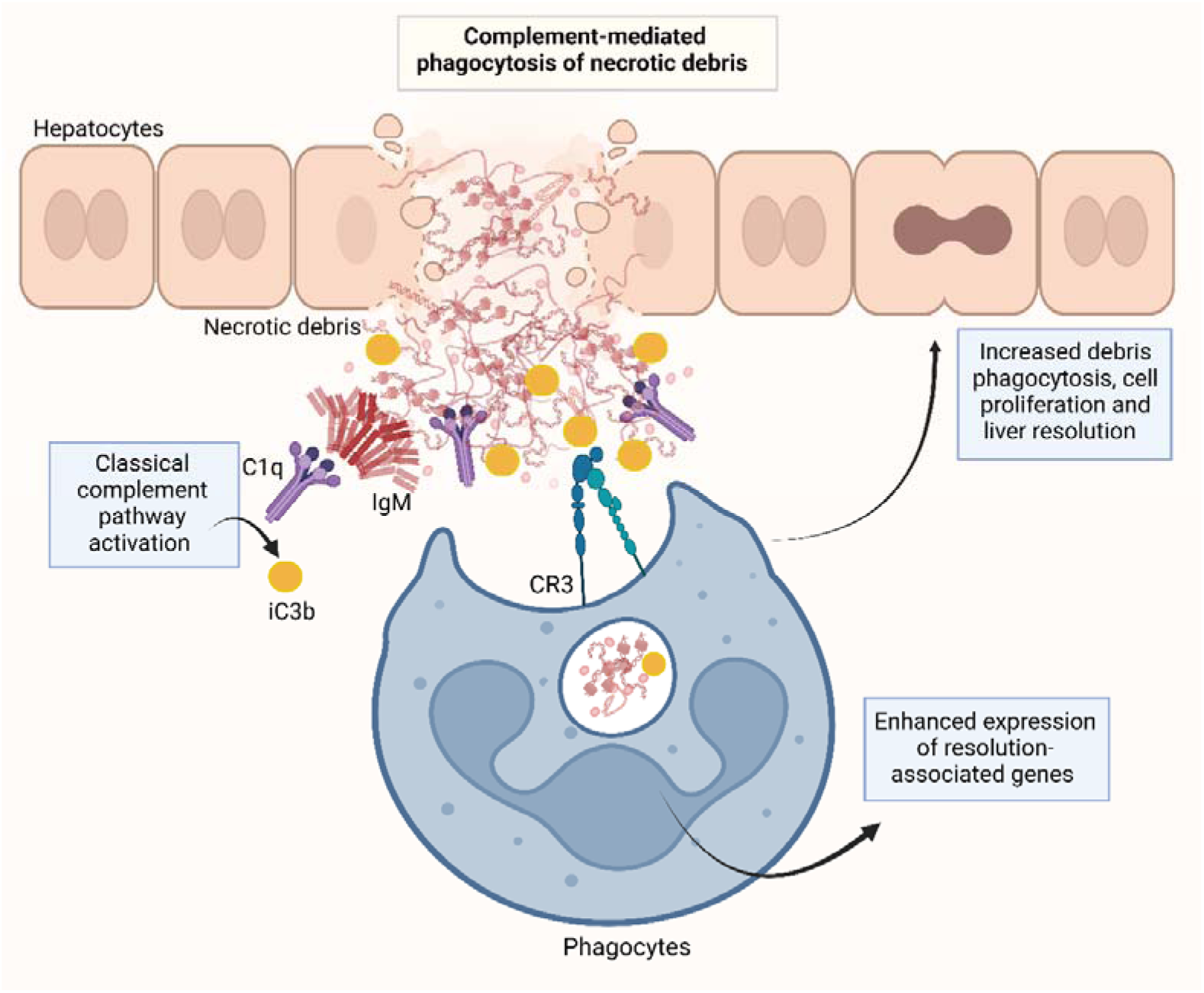

**Key points:**

- The complement cascade is activated on necrotic cell debris *in vivo* via the classical pathway
- Deficiency in complement C3 impairs necrotic debris clearance and liver recovery after injury
- Complement-mediated debris clearance is performed by neutrophils, monocytes and macrophages
- Human neutrophils depend on complement opsonization to phagocytose necrotic cell debris efficiently

## Introduction

The complement cascade is an essential part of the innate immune system, comprising over 50 soluble and membrane-bound proteins that work together to destroy pathogens and maintain tissue homeostasis by removing dying cells.^1^ It involves three distinct pathways: the classical, alternative and lectin pathway. All rely on different molecules for initial cascade activation, yet they converge to a central step where the C3 convertase cleaves complement component C3 into the anaphylatoxin C3a and the opsonin (i)C3b. Invading microorganisms and dead cells become opsonized by (i)C3b, which enables phagocytic cells to recognize and internalize them. Depending on the nature of the dying cell, diverse molecules may facilitate clearance. For instance, apoptotic bodies unveil ‘eat me’ signals on their membranes, in particular phosphatidylserine, which is recognized by a multitude of phagocytic receptors on leukocytes.^2^ Complement also contributes to this removal via binding of C1q to the apoptotic cell, thereby opsonizing it and activating the complement pathway to facilitate clearance through phagocytosis.^3,4^ Even though the general role of (i)C3b in phagocytosis is well-established, so far nearly all research on complement-mediated phagocytosis focused on microorganism clearance and efferocytosis (of apoptotic cells).^5–8^ The removal of cellular corpses resulting from necrosis, referred to as necrotic cell debris, has been largely overlooked both in terms of mechanism description and physiological impact *in vivo*.

Necrotic cell death diverges morphologically and immunologically from apoptosis due to plasma membrane rupture, the spilling of intracellular contents into the surrounding tissue, and the subsequent inflammatory response.^9^ These contents serve as damage-associated molecular patterns (DAMPs), which include ATP^10^, high-mobility group box 1 (HMGB1)^11^, actin^12^, mitochondria-derived molecules^13^ and DNA^14^, among others, and will interact with cognate pattern-recognition receptors (PRRs) on immune cells. For example, during drug-induced liver injury, widespread hepatocyte necrosis results in substantial DNA release from necrotic cells and causes intense TLR9-dependent inflammation.^15,16^ Also, HMGB1 interaction with the receptor for advanced glycation endproducts (RAGE) triggers neutrophil recruitment to the site of necrosis, contributing to injury amplification.^17^ Other DAMPs such as histones and F-actin released from necrotic cells also contribute heavily to immune responses through Clec2d and Clec9a recognition, respectively.^18,19^ These highlight the importance of limiting the accumulation of debris/DAMPs in conditions where necrosis is prominent, such as drug-induced liver injury, atherosclerosis, stroke, severe trauma, and burn injuries.

Phagocytosis is a cellular process of recognition and ingestion of particles larger than 0.5 µm, which promotes tissue homeostasis and elimination of microorganisms. Phagocytes recognize targets through specialized surface receptors, including non-opsonic receptors such as Dectin-1, Mincle, CD14 and CD36, which detect conserved molecular patterns, as well as various opsonic receptors. Complement receptors [*e.g.* CR1, CR3 (CD11b/CD18), CR4] and Fcγ receptors (*e.g.* FcγRI, FcγRIII, FcγRIV) are typical opsonic receptors recognizing particles bound by complement opsonins and antibodies, respectively.^20^ In general, complement has been implicated in the processing and removal of self-antigens, since the clearance of apoptotic cells *in vitro* by phagocytes was shown to be dependent on opsonization by C1q and C3.^21,22^ This central role is also observed in humans by the increased susceptibility to autoimmune disorders such as systemic lupus erythematosus (SLE) and rheumatoid arthritis in individuals with complement deficiencies.^23^ As an example, around 90% of patients with a C1q deficiency present SLE-like autoimmune manifestations due to the inappropriate clearance of immune complexes and apoptotic cells.^24^ Even though the role of complement in the clearance of apoptotic cells is clear, its contribution to the clearance of necrotic cells is poorly understood, with *in vivo* evidence lacking. In addition, the phagocyte populations that might be involved in necrotic debris removal and the impact of such clearance remain unclear.

Our group has recently shown that natural IgM and IgG antibodies are essential for the clearance of necrotic debris *in vivo*.^25^ Considering the substantial capacity of antibodies to initiate complement, the contribution of complement activation to necrotic cell debris clearance may be central. Therefore, we investigated complement activation in response to necrotic injury in mouse models of drug-induced liver injury and focal thermal injury (FTI) of the liver. We used intravital microscopy (IVM) to unveil the participation of complement in the clearance of necrotic debris *in vivo* and assessed its impact on the recovery from liver injury.

## Materials and methods

### Mice

8-12 weeks old male and female C57BL/6J and C57BL/6NRj mice were purchased from Janvier Labs. RAG2^-/-^ mice (C57BL/6N-Rag2Tm1/CipheRj) were bred in specific pathogen-free (SPF) conditions at the Animal Facility of the Rega Institute (KU Leuven). C3^-/-^ mice (B6.129S4-C3tm1Crr/J) and Itgam^-/-^ mice (B6.129S4-Itgamtm1Myd/J) were purchased from The Jackson Laboratory. Mice were housed in acrylic filtertop cages with an enriched environment (bedding, toys and small houses). Mice were kept under a controlled light/dark cycle (12/12h) at 21 °C with water and food provided *ad libitum*. All experiments were approved and performed following the guidelines of the Animal Ethics Committee from KU Leuven (P125/2019).

### APAP-induced liver injury model

Mice were starved for 15h and given a single oral gavage of 600 mg/kg APAP (Sigma-Aldrich) dissolved in warm PBS. After 24, 48 or 72h, mice were sacrificed under anesthesia containing 80 mg/kg ketamine and 4 mg/kg xylazine, whereafter liver and blood were harvested. ALT in serum was determined with a kinetic enzymatic kit (Infinity, Thermo Fisher Scientific) according to the manufacturer’s instructions.

### C3 and C3a ELISA

Serum levels of C3a were measured using an enzyme-linked immunosorbent assay (ELISA). A 96-well plate (Corning) was coated with 2 µg/ml rat anti-mouse C3a (clone I87-1162, cat 558250, BD Biosciences) in PBS and incubated at 4°C overnight. The plate was blocked in blocking buffer (PBS with 0.1% casein and 0.05% Tween20) for 2h at room temperature (RT). Purified mouse C3a (cat 558618, BD Biosciences) was used as standard ranging from 50 ng/ml to 0.39 ng/ml in blocking buffer. Samples were diluted (1:1000) in blocking buffer and incubated for at least 2h at RT. After washing, 0.5 µg/ml biotinylated rat anti-mouse C3a (clone I87-1162, cat 558251, BD Biosciences) in blocking buffer was added and incubated for 1h at RT. Next, streptavidin-horseradish peroxidase (R&D Systems) was added in blocking buffer and incubated for 30 min at RT, followed by adding a 3,31,5,51-tetramethylbenzidine (TMB) solution. The reaction was stopped by adding 1M H_2_SO_4_ and the absorbance was measured at 450 nm using a PowerWave XS plate reader (BioTek Instruments). Serum levels of mouse C3 were determined by a commercially available C3 ELISA kit (ab157711, Abcam) according to the manufacturer’s instructions.

### Neutrophil purification

Human neutrophils were purified from whole blood of healthy volunteers by immunomagnetic negative selection (EasySep^TM^ Direct Human Neutrophil Isolation Kit, StemCell Technologies, Vancouver, Canada) according to the manufacturer’s instructions. Ethical permission for use of human blood-derived leukocytes was obtained with the ethical committee from University Hospital Leuven (UZ Leuven, S58418). Mouse bone marrow neutrophils were extracted from femurs and tibias of C57BL/6J mice by flushing the bones with 5 mL cold RPMI-1640 medium using a 26-gauge needle. Cells were filtered through a 70 µm nylon strainer and further purified with the EasySep^TM^ mouse neutrophil enrichment kit (StemCell Technologies), following the manufacturer’s instructions.

### Histopathology

Liver sections were stained with hematoxylin and eosin (H&E) and used to estimate hepatic necrosis via measurement of the necrotic area in the images. The livers were washed with 0.9% NaCl and fixed in 4% buffered formalin. Subsequently, the samples were dehydrated in ethanol solutions, bathed in xylol and included in histological paraffin blocks. Tissue sections of 5 μm were obtained using a microtome and stained with H&E. Sections were visualized using a BX41 optical microscope (Olympus) and images were obtained using the Moticam 2500 camera (Motic) and Motic Image Plus 2.0ML software.

### Cryosection immunostaining for C3b, C1q, Ki67, fibrin(ogen) and IgM

The left liver lobes of mice were harvested, embedded in Tissue-Tek O.C.T. Compound (Sakura Finetek Europe) and snap frozen in liquid nitrogen. 10 µm sections were cut using a Cryostat Microm CryoStar and subsequently fixed, permeabilized and blocked. Sections were incubated overnight at 4°C with 10 µg/ml polyclonal rabbit anti-human/mouse fibrin(ogen) (Dako), 5 µg/ml rat anti-mouse C3b/iC3b (clone 3/26, Hycult Biotec) and 5 µg/ml rabbit anti-mouse C1q (clone 4.8, Abcam). Secondary antibodies were added for 3h at RT: Alexa Fluor 647 donkey anti-rabbit, Rhodamine RED-X (RRX) donkey anti-mouse IgM, Alexa Fluor 488 donkey anti-rat, Alexa Fluor 560 donkey anti-rabbit (all at 10 µg/mL, Jackson ImmunoResearch). 10 µg/ml of Hoechst was added for 30 min at RT to stain nuclei. Finally, slides were mounted with ProLong Diamond Antifade Mountant. Images were captured using the Andor Dragonfly High-Speed Confocal Microscope (Oxford Instruments) or a Zeiss Axiovert 200M fluorescence microscope, and analyzed with FIJI. 8 images were acquired per liver with a 25X objective. Stained areas were selected using the threshold tool in FIJI, from which the percentage area of staining was determined. Pearson’s coefficient was calculated in FIJI using the JACoP plugin. Comparisons between WT and Rag2^-/-^ mice were normalized to the degree of injury [% of fibrin(ogen) labeling].

### Confocal Intravital microscopy

Mice were anaesthetized with a subcutaneous injection of 80 mg/kg ketamine and 4 mg/kg xylazine. For the experiments with APAP-induced liver injury, fluorescent antibodies (4 µg/mouse) and dyes (2 µl of a 10 mM Sytox Green solution; Thermo Fisher Scientific) were dissolved in 100 µl sterile PBS and injected intravenously 10 minutes before the surgery. The surgical procedure is described in detail in Marques *et al*.^26^ For the FTI experiments, 1 mm^3^ burns were made with a cauterizer and the injury site was then stained with 10 µl of pHrodo Red succinimidyl ester (SE) (4 µM; Thermo Fisher Scientific). The incision was stitched, and after 6h, mice were again anaesthetized with ketamine and xylazine for imaging of the burn site by IVM. Images were taken every 30 sec for at least 30 min with the Dragonfly Spinning-Disk Confocal Microscope (Oxford Instruments). Quantification of phagocytosis was done in a blind manner and counted manually. The percentage of Sytox Green labeling was determined from 2 mosaic images, each composed of 16 images per mouse, with FIJI software using thresholding. The % of CD11b+ cells containing DNA was determined using Imaris software. Surfaces overlaying live cells and DNA debris were generated for 3D images and counted manually.

### In vitro phagocytosis

Necrotic debris was generated from HepG2 cells by inducing mechanical disruption with a pellet mixer for 5 min in 0.1 M sodium bicarbonate (pH 8.5). The necrotic debris was labeled by adding 2 µl of 10 mM pHrodo Red SE (Thermo Fisher Scientific) solution per 10×10^6^ cells. The debris was opsonized with 20% normal human serum, C1q^-/-^ serum (Complement Technology) or C3^-/-^ serum (Complement Technology) in PBS for 1h at 37 °C. Opsonized debris was added to purified neutrophils in a 1:10 cell/debris ratio. Neutrophils were stimulated with 10^-7^ M N-formyl-Met-Leu-Phe (fMLF; Sigma-Aldrich) for human neutrophils or 1 µM WKYMVM (Phoenix Pharmaceuticals) for mouse neutrophils, and labeled with 1 µM calcein AM viability dye (Invitrogen). Two 3D mosaics are captured per well, each comprising 9 overlapping images taken after 3h incubation at 37°C with the 25X objective of the Dragonfly Confocal Microscope (Oxford Instruments). Each condition was plated in duplicate and replicated at least 3 times. 3D reconstructions were generated using Imaris software. Additionally, with Imaris, surfaces were overlaid onto live cells and necrotic debris through thresholding, after which the volume of overlap was calculated in µm^3^.

### Flow cytometry

Liver lobes were surgically removed, put in MACS tubes with RPMI-1640 (Biowest Riverside) and minced with a gentleMACS Dissociator (Miltenyi Biotec). To this suspension, 2.5 mg collagenase D (Roche) and 1 mg DNAse I were added per liver for 1h at 37°C. The cell suspension was washed with PBS (300 g, 5 min, 4 °C). Non-parenchymal cells were separated by density gradient centrifugation at 60 g for 3 min at 4 °C. Supernatant was collected and filtered through a 70 µm nylon cell strainer. After centrifugation (300 g, 5 min, 4°C), ACK Lysing buffer (Gibco) was added for 10 min to the pellet to lyse red blood cells. 1×10^6^ cells were collected in FACS tubes and washed with PBS (300 g, 5 min, 4 °C). Zombie Aqua Fixable Viability dye (Biolegend) together with mouse FcR blocking Reagent (Miltenyi Biotec) were incubated for 15 min in the dark. Then, cells were washed with PBS supplemented with 0.5% bovine serum albumin (BSA) and 2 mM EDTA and the fluorescently labeled antibodies were incubated for 25 min at 4°C in the dark. After a final washing step, cells were read in a Fortessa X20 (BD Biosciences). Data was analyzed using FlowJo 10.8.1 software.

### RNA extraction from human neutrophils fed necrotic debris

HepG2 cells were mechanically lysed using a 22G syringe in PBS with 20 µg/ml RNAse A (Sigma-Aldrich) to generate RNA-free necrotic debris. The debris was incubated for 30 min at 37°C to allow enzyme activity. Then, the debris was washed twice with PBS containing 2 mM EDTA and 0.1% BSA, and centrifuged at 60g for 3 min to remove intact cells. The debris was opsonized with 20% serum or C3-depleted serum (Complement Technology) and then incubated for 1h at 37°C. Neutrophils from healthy donors were purified by immunomagnetic negative selection with an EasySep kit and stimulated with 10^-7^ M fMLF. Cells and debris were co-incubated in a 6 well plate at a 1:10 cell/debris ratio and centrifuged at 300g for 5 min before being incubated for 3h at 37°C. Cells were harvested and total RNA was extracted by lysing the cells with β-mercaptoethanol and a Rneasy Plus Mini Kit (Qiagen) following the manufacturer’s instructions. After extraction, total RNA quality and quantity was determined using a Nanodrop.

### qPCR

cDNA was obtained by reverse transcription using the high-capacity cDNA Reverse Transcriptase kit (Applied Biosystems). mRNA levels were analyzed by quantitative PCR using a TaqMan Gene Expression Master Mix (Applied Biosystems) and a 7500 Real-Time PCR System apparatus. Expression levels of genes of interest were normalized for the average RNA expression of three housekeeping genes (CDKN1A, 18s and GAPDH) using the 2^−ΔΔCT^ method.^27^

### Statistical analysis

Data were analyzed using GraphPad Prism v9.3.1. All data are expressed as mean ± standard error of the mean (SEM). A Shapiro-Wilkinson test was performed to check for normality. Normally distributed data were analyzed with a Student’s t test or One-way ANOVA. Non-parametric data were analyzed with a Mann-Whitney test or Kruskal-Wallis test. Grubb’s test (extreme studentized deviate) was applied to determine significant outliers. A p-value equal or lower than 0.05 was considered significant.

## Results

### Complement proteins C1q and (i)C3b are deposited at sites of necrotic liver injury

To assess complement activation at necrotic injury sites, a mouse model of paracetamol/acetaminophen (APAP)-induced liver injury was used, as it is characterized by extensive death of hepatocytes through necrosis. A sublethal dose of 600 mg/kg APAP was administered via oral gavage, causing significant liver damage as early as 12h after administration. In this acute model, hepatocellular necrosis was determined by elevated levels of serum ALT (**supplemental Figure 1A**), an enzyme primarily found in hepatocytes that serves as a biomarker for liver damage. Using histopathology, necrotic lesions were observed around the centrilobular veins, a typical pattern for APAP-induced injury, with the highest severity 12h after APAP administration (**supplemental Figure 1B-C**). At 48h, the injury decreased significantly, as depicted by lower serum ALT levels and reduced necrotic areas (**supplemental Figure 1A-B**). Fibrin deposition, known for its specific accumulation at necrotic sites after the activation of the coagulation cascade^28^, was evaluated over time on liver cryosections to estimate the area of necrosis. Peak staining was observed after 24h, whereafter it gradually decreased (**Figure 1A, C; supplemental Figure 2A**). A similar pattern was observed for IgM, with the highest deposition occurring after 24h and diminishing at later timepoints (**Figure 1B and D; supplemental Figure 2B**). Remarkably, C1q and (i)C3b deposition were maximal mostly after 48h (**Figure 1E-F**), indicating that the deposition of antibodies precedes complement activation through C1q binding and C3 cleavage. Pearson’s correlation coefficient between (i)C3b and fibrin was significantly higher in comparison to (i)C3b and intact cell nuclei in the liver (**Figure 1G**). In addition, the other components of the classical complement pathway, IgM and C1q, also had high colocalization with (i)C3b (**Figure 1H**). All this shows that complement proteins are deposited specifically at sites of necrotic injury in the liver. These data were supported by significantly lower C3 and C3a levels in the serum of APAP-treated mice (**Figure 1I-J**), which confirm complement activation and anaphylatoxin consumption in response to injury.

**Figure 1.**
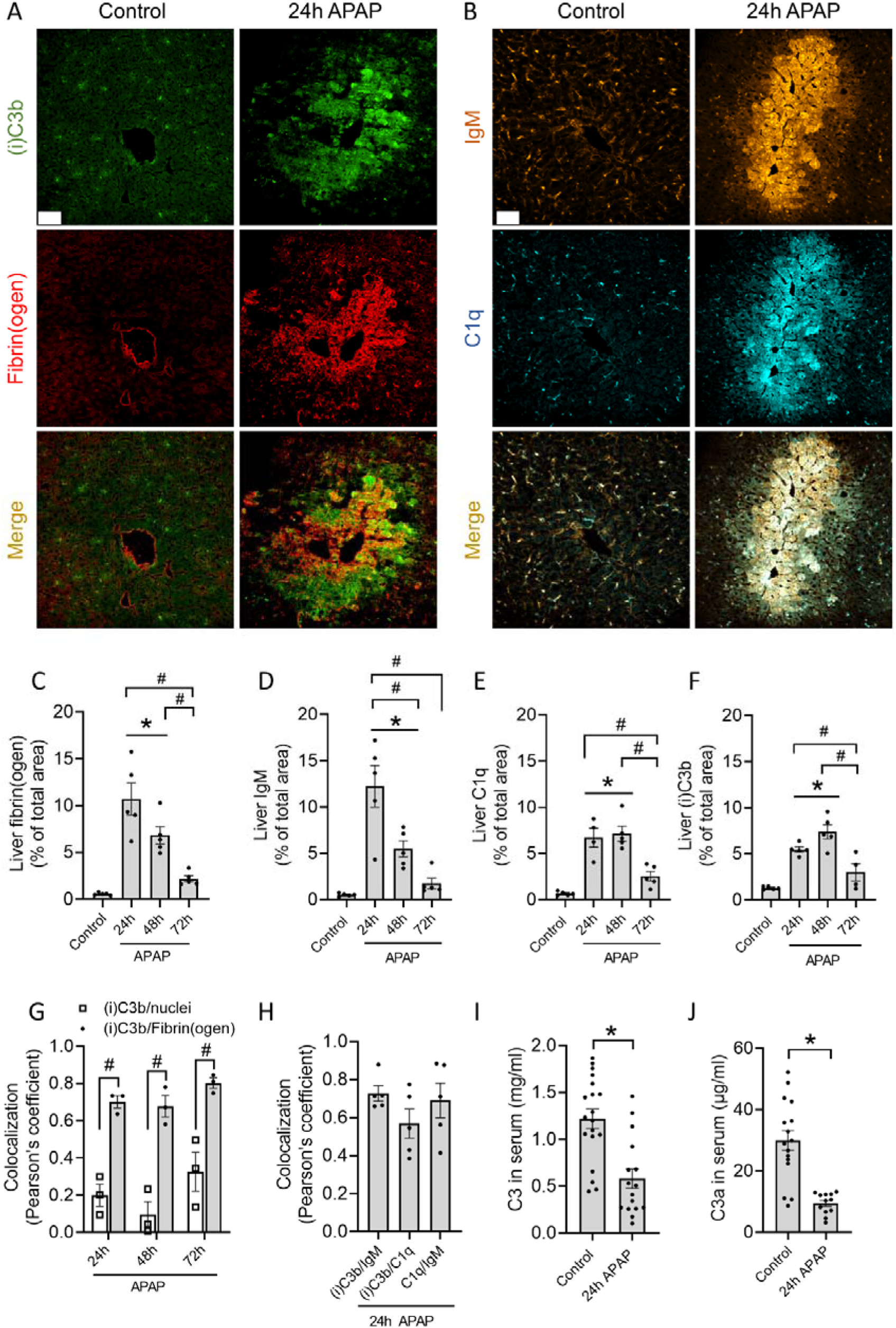
Complement proteins C1q and (i)C3b are deposited at sites of necrotic liver injury. (A-B) Representative immunofluorescence images of liver cryosections from control mice or mice challenged with acetaminophen (APAP; 600 mg/kg) for 24h. Green: (i)C3b; red: fibrin(ogen); Orange: IgM; Cyan: C1q. Scale bar represents 50 µm. (C-F) Quantification area percentage stained with fibrin(ogen) (C), IgM (D), C1q (E) and (i)C3b (F) in liver cryosections of control mice and mice receiving an APAP overdose for 24, 48 or 72h. (G) Pearson’s correlation coefficient of (i)C3b and nuclei (Hoechst) or fibrin(ogen) in the injured livers 24, 48 and 72h after APAP overdose. (H) Pearson’s correlation coefficient between (i)C3b, IgM and C1q in the injured livers 24h after APAP overdose. (I) C3 levels in the serum of control mice or challenged with APAP for 24h. (J) C3a levels in the serum of control mice or challenged with APAP for 24h. Image quantifications were pooled from 8 fields of view. Each dot represents a single mouse. Data are represented as mean ± SEM. *p≤0.05 compared to control; #p≤0.05 between indicated groups. APAP = acetaminophen.

### Classical complement pathway is activated in response to necrotic debris

To investigate how (i)C3b opsonization of necrotic debris was initiated, Rag2^-/-^ mice were used. In these mice, the development of mature T and B cells is stopped at the pro-T and B cell stages, preventing the production of antibodies. Activation of the classical complement pathway requires the association of C1q with target-bound IgM or multiple IgGs, thus in these Rag2^-/-^ mice, this complement pathway cannot be activated. First, the absence of IgM in these mice was confirmed by immunostaining, which revealed that IgM labeling in the injured liver was essentially absent (**Figure 2A-C**). Interestingly, 24h after APAP overdose, liver cryosections showed significantly increased fibrin deposition in Rag2^-/-^ mice compared to WT mice (**Figure 2D**). Because the degree of necrotic injury also affects the degree of C1q and (i)C3b deposition, complement immunostaining was normalized to the area of fibrin staining. With this approach, we observed no difference in C1q binding to necrotic areas in Rag2^-/-^ mice compared to WT (**Figure 2E**). Similarly, the mean fluorescence intensity (MFI) of C1q was not different between WT and Rag2^-/-^ mice, indicating that C1q binding does not rely on antibody opsonization of necrotic debris (**Figure 2F**). However, the area of (i)C3b deposition was significantly decreased in Rag2^-/-^ mice, with also significantly lower MFI in (i)C3b-stained areas (**Figure 2G-H**), even though it has been shown that Rag2^-/-^ mice have elevated C3 levels in blood.^29^ These data indicate that loss of IgM and IgG antibodies diminishes the level of complement activation on necrotic sites, even though C1q binding remains unaffected. These data also demonstrate that the activation of the classical pathway in response to necrotic debris is responsible for the majority of (i)C3b deposition in injury sites.

**Figure 2.**
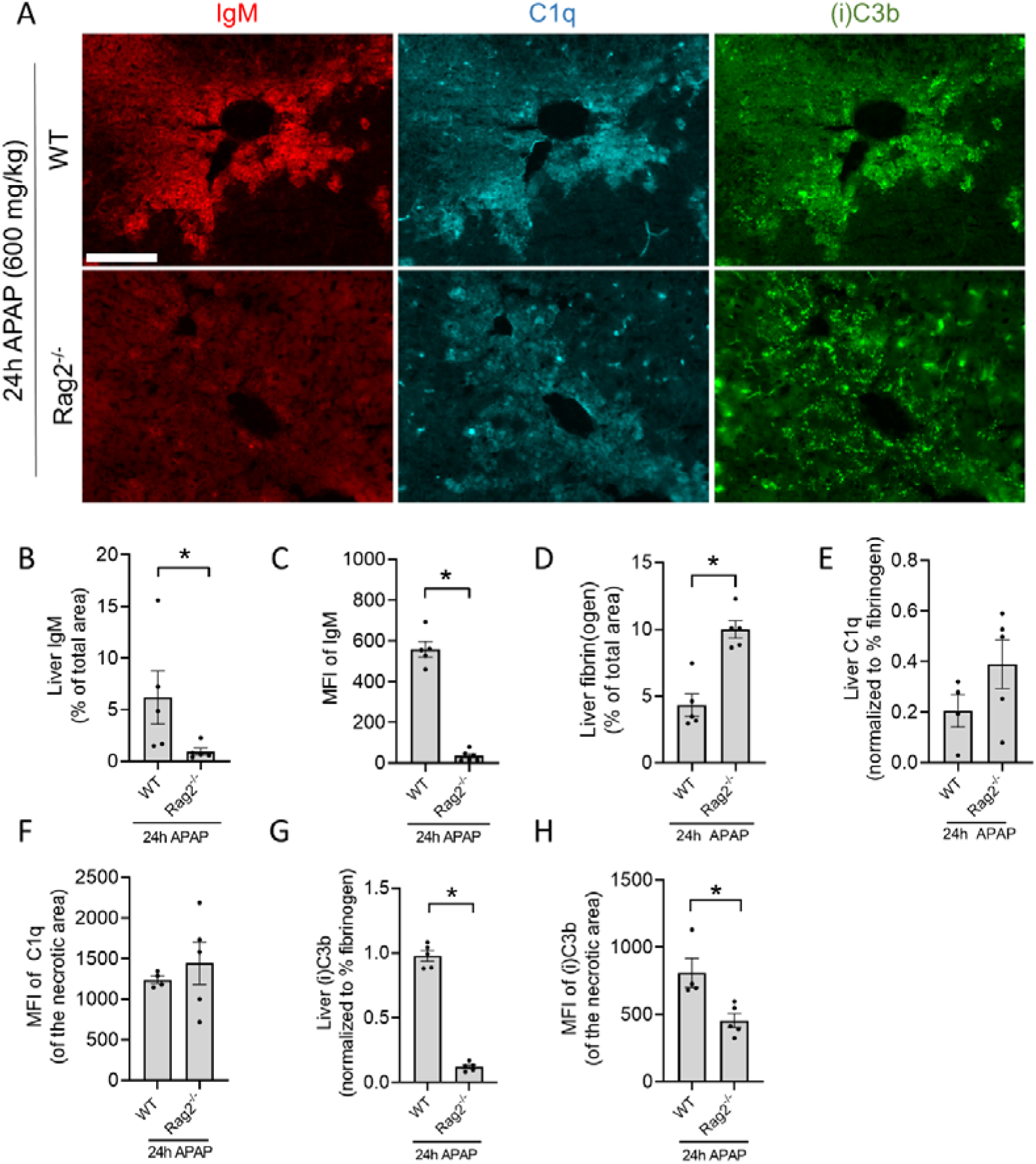
The classical complement pathway is activated in response to necrotic debris. (A) Representative immunofluorescence images of liver cryosections from WT and Rag2^-/-^ mice 24h after an overdose of acetaminophen (APAP; 600 mg/kg). Red: IgM; Cyan: C1q; Green: (i)C3b. Scale bar represents 100 µm. (B) Quantification of area percentage stained with IgM in liver cryosections of WT and Rag2^-/-^ mice 24h after APAP overdose. (C) Mean fluorescence intensity (MFI) of IgM staining at the necrotic injury sites in liver cryosections. (D) Quantification of area percentage stained with fibrin(ogen) in liver cryosections of WT and Rag2^-/-^ mice 24h APAP overdose. (E) Quantification of C1q deposition in the liver of WT and Rag2^-/-^ mice 24h after APAP overdose, normalized to the degree of fibrin(ogen) staining. (F) Mean fluorescence intensity of C1q staining at the necrotic injury sites in liver cryosections. (G) Quantification of (i)C3b deposition in the liver of WT and Rag2^-/-^ mice 24h after APAP overdose, normalized to the degree of fibrin(ogen) staining. (H) Mean fluorescence intensity of (i)C3b staining at the necrotic injury site of liver cryosections. Image quantifications were pooled from 8 fields of view. Each dot represents a single mouse. Data are represented as mean ± SEM. *p≤0.05 compared to WT mice. APAP= acetaminophen, MFI = Mean fluorescence intensity.

### C3**^-/-^** mice have delayed recovery from liver injury

To investigate the contribution of C3 to the resolution of necrotic liver damage, C3^-/-^ mice were subjected to APAP overdose and evaluated at 2 timepoints: a) after 24h, to assess the peak of injury and b) after 48h, to observe the degree of tissue repair. No differences in serum ALT levels were observed after 24h, while significantly higher levels of ALT were found after 48h in C3^-/-^ mice compared to WT mice (**Figure 3A**). In addition, fibrin staining of liver cryosections revealed no differences at the peak of injury, whereas more fibrin staining was found in C3^-/-^ mice at the later timepoint (**Figure 3B**), indicating that C3 deficiency leads to larger, unresolving necrotic areas in the liver. These data were corroborated by the larger necrotic areas also found in Rag2^-/-^ mice, which presented a defect in complement activation in the liver (**Figure 2D**). During the liver regeneration phase, cellular proliferation can be estimated by the expression of Ki67 in liver cryosections. Using this approach, we observed a significant decrease in cell proliferation in C3^-/-^ mice 48h after APAP overdose (**Figure 3C**), confirming that the absence of C3 impairs liver regeneration and recovery from injury.

**Figure 3.**
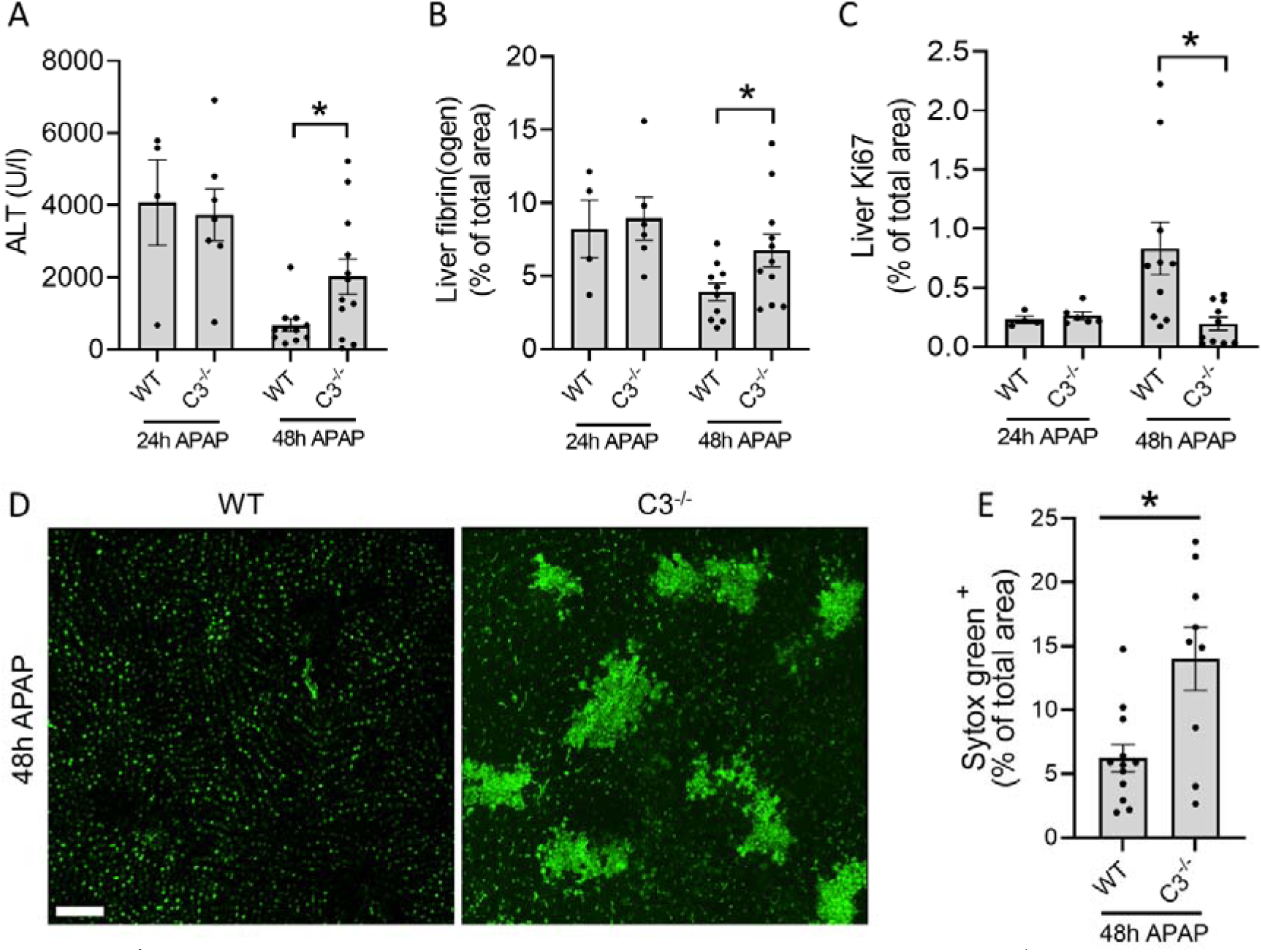
C3^-/-^ mice have delayed recovery from liver injury. (A) Serum ALT levels in WT and C3^-/-^ mice 24 and 48 hours after APAP overdose. (B) Quantification of area fraction stained with fibrin(ogen) in liver cryosections. (C) Quantification of area fraction stained with Ki67 in liver cryosections of APAP-challenged WT and C3^-/-^ mice. (D) Representative IVM images of WT and C3 mice showing the deposition of DNA (Sytox Green) in the liver 48 hours after APAP overdose (600 mg/kg). Scale bar represents 200 µm. (E) Quantification of Sytox Green labeling in the liver of APAP-challenged WT and C3^-/-^ mice after 48h. Images were analyzed with Imaris software. Each dot represents a single mouse. Data are represented as mean ± SEM. *p≤0.05 between indicated groups. APAP= acetaminophen, ALT = alanine aminotransferase.

In order to assess directly if the amount of necrotic debris in injured tissues would also be impacted by C3 deficiency, we performed confocal IVM of mouse livers. Considering that DNA is abundantly released by hepatocytes^16^, we chose to measure the amount of DNA exposed in the liver 48h after the APAP challenge using the membrane-impermeable DNA dye Sytox Green. Interestingly, WT mice presented minimal extracellular DNA in the liver at the 48h timepoint, which is consistent with the removal of necrotic debris and tissue recovery at that phase (**Figure 3D-E**). Moreover, the vast majority of the fluorescent signal observed in the images of WT mice consisted of background fluorescence from healthy hepatocyte nuclei (**Figure 3D**). However, in C3^-/-^ mice, we observed a significantly larger area still covered by extracellular DNA debris, demonstrating that these mice have a clear defect in the removal of necrotic debris from the liver (**Figure 3D-E**). These data show that the poor recovery from liver injury in C3^-/-^ mice is connected to an impairment in necrotic cell debris clearance.

### Phagocytosis of necrotic DNA debris depends on complement activation

To explore how debris persisted in injury sites, an analysis of the recruited leukocytes and their ability to take up debris was performed. The inflammatory response triggered by liver injury led to the recruitment of CD11b^+^ leukocytes to necrotic areas identified by the extracellular DNA staining (**Figure 4A, upper panels**). These cells consisted primarily of inflammatory monocytes (CCR2^+^), neutrophils (Ly6G^+^) and macrophages (F4/80^+^) (**Figure 4A, lower panels; supplemental Figure 3A -C**). Of interest, CD11b, the α_M_ subunit of the complement receptor CR3 (CD11b/CD18), which is known for its involvement in complement-mediated phagocytosis, was increased in neutrophils and monocytes during liver injury **(Supplemental Figure 3D-F)**. We then inquired whether CD11b^+^ cells were able to internalize extracellular DNA debris using IVM.

**Figure 4.**
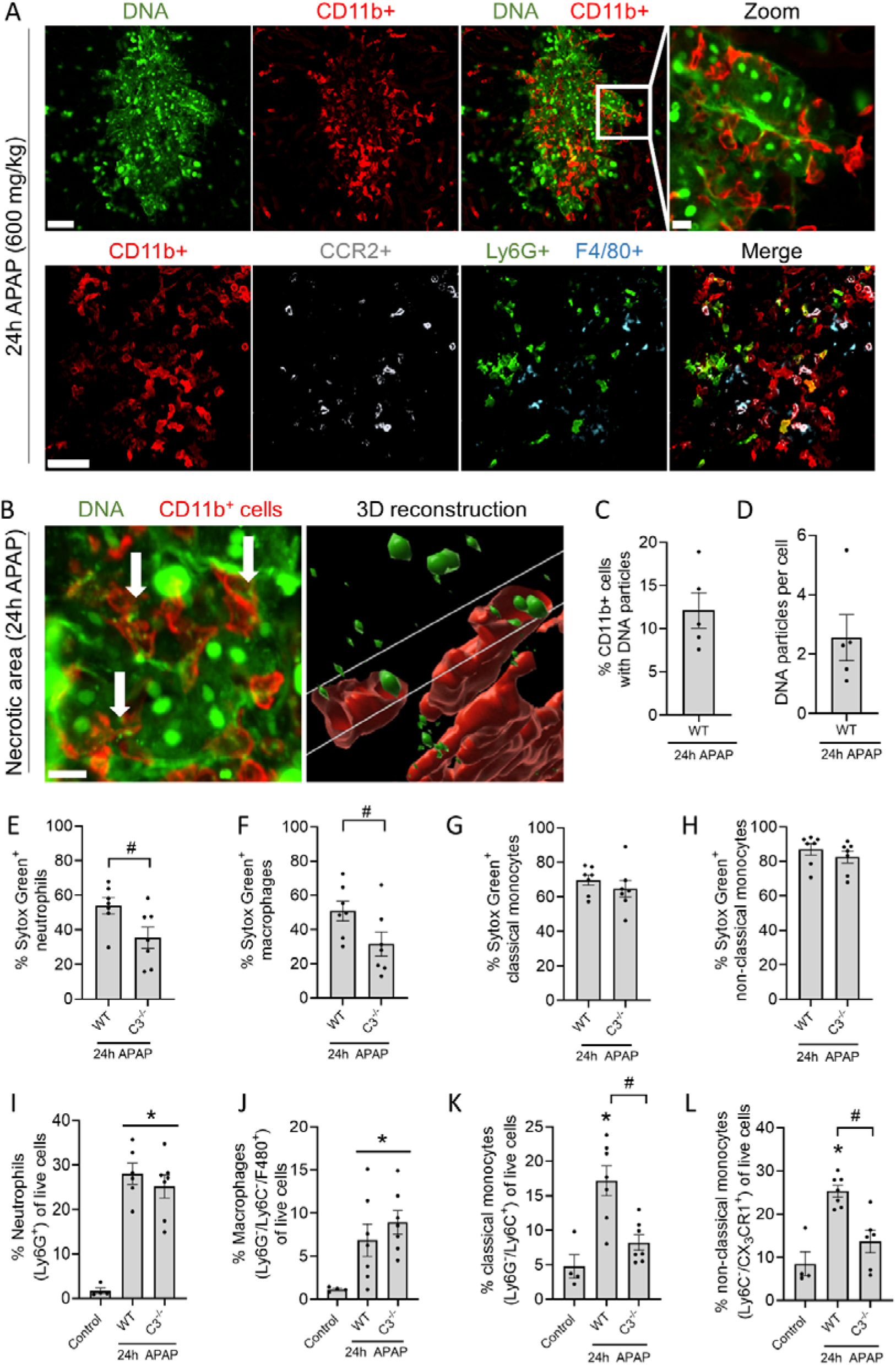
Phagocytosis of necrotic DNA debris depends on complement activation. (A) Representative IVM images of the injured liver 24h after APAP overdose. Sytox Green (Green; DNA), CD11b (Red; leukocytes), CCR2 (White; monocytes), Ly6G (Green; neutrophils) or F4/80 (Cyan; macrophages). Upper and lower panels are from different experiments. Scale bar represents 50 µm, scale bar zoomed image represents 10 µm. (B) Representative IVM image of the injured liver 24h after APAP and an image of a 3D reconstruction with Imaris software. 2 µl of the DNA impermeable dye Sytox Green was injected 1h before imaging. Leukocytes were labeled with anti-CD11b antibody (Red). Scale bar represents 50 µm. (C) Percentage of CD11b cells containing DNA particles in the injured liver of mice quantified by IVM. (D) Average number of DNA particles per CD11b cell in the injured liver of mice quantified by IVM. (E-L) Flow cytometry of non-parenchymal cells identifying the percentage of neutrophils (Ly6G^+^), non-classical monocytes (Ly6C^-^ / CX_3_CR1^+^), classical monocytes (Ly6G / Ly6C) and macrophages (Ly6G /Ly6C / F4/80) in the injured liver and the percentage of those leukocytes containing extracellular DNA (Sytox Green). The cell-impermeable DNA dye Sytox Green was injected 2h before harvesting the liver of control, WT and C3^-/-^ mice that received an APAP overdose 24h prior. Data are represented as mean ± SEM. Each dot represents a single mouse. *p≤0.05 compared to control; #p≤0.05 between indicated groups. APAP = acetaminophen.

Numerous CD11b^+^ leukocytes were visualized deep within necrotic areas using Z-stacks, and multiple cells contained Sytox Green^+^ phagosomes (**Figure 4B and supplemental video 1**). In total, 12% of the CD11b^+^ cells contained DNA particles, with an average of 2 particles per cell (**Figure 4C-D**). Internalization of DNA debris was confirmed upon intravenous administration of DNAse to remove the bulk of extracellular DNA in necrotic areas. CD11b^+^ leukocytes with DNA-containing vesicles remained after DNAse injection, indicating that the DNA particles were located intracellularly, likely in phagosomes, which are not accessible to the circulating DNAse treatment (**Supplemental Figure 3G and supplemental video 2**).

Following the intravital imaging of DNA debris uptake and the observation of impaired DNA removal in the absence of C3 (**Figure 3D-E**), flow cytometry was performed to quantify DNA uptake in WT and C3^-/-^ mice. Again, the membrane-impermeable DNA dye Sytox Green was injected intravenously 2h before sacrificing mice that received an APAP overdose 24h prior. This allowed sufficient time for the fluorescently-labeled necrotic DNA to be phagocytosed. We observed that a significantly lower percentage of neutrophils and macrophages were able to internalize DNA debris in the absence of C3, demonstrating that debris clearance depends at least partially on complement opsonization (**Figure 4E-F**). Importantly, no differences were observed in the number of neutrophils (Ly6G^+^) and macrophages (Ly6G^-^ / Ly6C^-^ / F4/80^+^) present in the injured liver between WT and C3^-/-^ mice (**Figure 4I-J**), suggesting that impaired DNA removal observed in C3^-/-^ mice is due to the reduced phagocytic capacity of these cells. In contrast, classical and non-classical monocytes did not require C3 to take up DNA debris in the injured liver (**Figure 4G-H**), suggesting that different leukocyte populations utilize distinct mechanisms for debris clearance. However, the number of classical (Ly6G^-^ / Ly6C^+^) and non-classical monocytes (Ly6C^-^ / CX_3_CR1^+^) recruited to the injured liver in C3^-/-^ mice was significantly reduced (**Figure 4K-L**), meaning that less monocytes reach the injured liver to perform debris phagocytosis. Overall, these data show that necrotic DNA debris is cleared by leukocytes in the liver and that neutrophils and macrophages require complement opsonization of debris for its uptake.

### Complement opsonization and clearance of necrotic cell debris are not exclusive to drug-induced liver injury

Reduced clearance of DNA debris in the absence of C3 was observed in a model of acute liver injury induced by APAP overdose. To validate whether this finding applies to other types of injuries, we investigated debris clearance in a model of focal thermal injury (FTI) of the liver. In the FTI model, necrotic lesions are induced locally with a hot needle, facilitating the observation of phagocytosis *in vivo*. This was challenging in the APAP model due to wide-spread necrosis throughout the liver causing an abundance of debris. The localized nature of the thermal injury allowed us to label necrotic debris by applying a droplet of the pH-sensitive dye pHRodo Red SE on top of the lesion. This dye binds covalently to proteins and exhibits increased fluorescence when the ingested material is processed in the acidic environment of a phagolysosome.

Our previous work demonstrated that neutrophils predominated in the injured area 6h after FTI, with monocytes being attracted after 12h.^25^ The entire image of the FTI shows distinct burn injury zones, with neutrophils mostly accumulating around the injury core (**supplemental Figure 4A**). Using IVM, neutrophils carrying pHRodo-labeled debris were observed crawling from the injury border into the necrotic core (**Figure 5A and supplemental video 3**). Approximately 75% of neutrophils at the injury site had pHRodo-containing phagosomes, whereas less than 5% of neutrophils in healthy areas of the liver phagocytosed debris (**Figure 5B**). This finding was confirmed by flow cytometry, which showed a significantly increased MFI of pHRodo in neutrophils and monocytes at the burn injury site compared to leukocytes in healthy areas (**supplemental Figure 4B-C**). Similarly to the APAP model, complement proteins C1q and (i)C3b were specifically deposited on necrotic lesions 6h after FTI (**Figure 5C**). Both components colocalized with each other and fibrin (**supplemental Figure 4D and E**). Quantification of necrotic debris clearance by flow cytometry demonstrated a significant decrease in the percentage of neutrophils and non-classical monocytes phagocytosing debris in C3^-/-^ mice compared to WT (**Figure 5D-E**). Interestingly, this phenomenon was not observed in classical monocytes nor macrophages (**Figure 5F-G**). To be noted, the number of neutrophils, macrophages and non-classical monocytes attracted to the injured liver did not differ between WT and C3^-/-^ mice, while the percentage of classical monocytes was significantly reduced in C3^-/-^ mice **(supplemental Figure 4F-I)**, showing that C3 deficiency affects the migration of classical monocytes to the injury, likely due to decreased C3a levels, rather than their capacity to phagocytose debris. Due to the upregulation of CD11b in response to liver injury **(supplemental Figure 3D-F)** and its known role in complement-mediated phagocytosis, the role of CR3 on debris uptake and liver resolution was investigated. In the FTI, using CD11b^-/-^ mice, we found a significant decrease in debris clearance by neutrophils and non-classical monocytes (**Figure 5D-E**). Again, this effect was not observed in classical monocytes and macrophages **(Figure 5F-G)**, corroborating our phagocytosis data using C3^-/-^ mice. Moreover, the phagocyte populations attracted to the injury site were not impaired by the deficiency in CD11b^-/-^, indicating that the defective clearance is not connected to inhibition of leukocyte recruitment **(supplemental Figure 4F-G)**. Conversely, no significant effect was observed on the progression of liver injury in CD11b^-/-^ mice or in WT mice that received a CD11b blocking antibody, as evidenced by similar ALT values and fibrin staining **(supplemental Figure 5A-E)** after APAP overdose. These data show that in the focal burn injury, neutrophils and monocytes migrate to the necrotic lesion to phagocytose necrotic debris, a process which depends on C3 and CD11b/CD18 for neutrophils and non-classical CX_3_CR1^+^ monocytes, whereas classical CCR2^+^ monocytes rely on other unidentified receptors.

**Figure 5.**
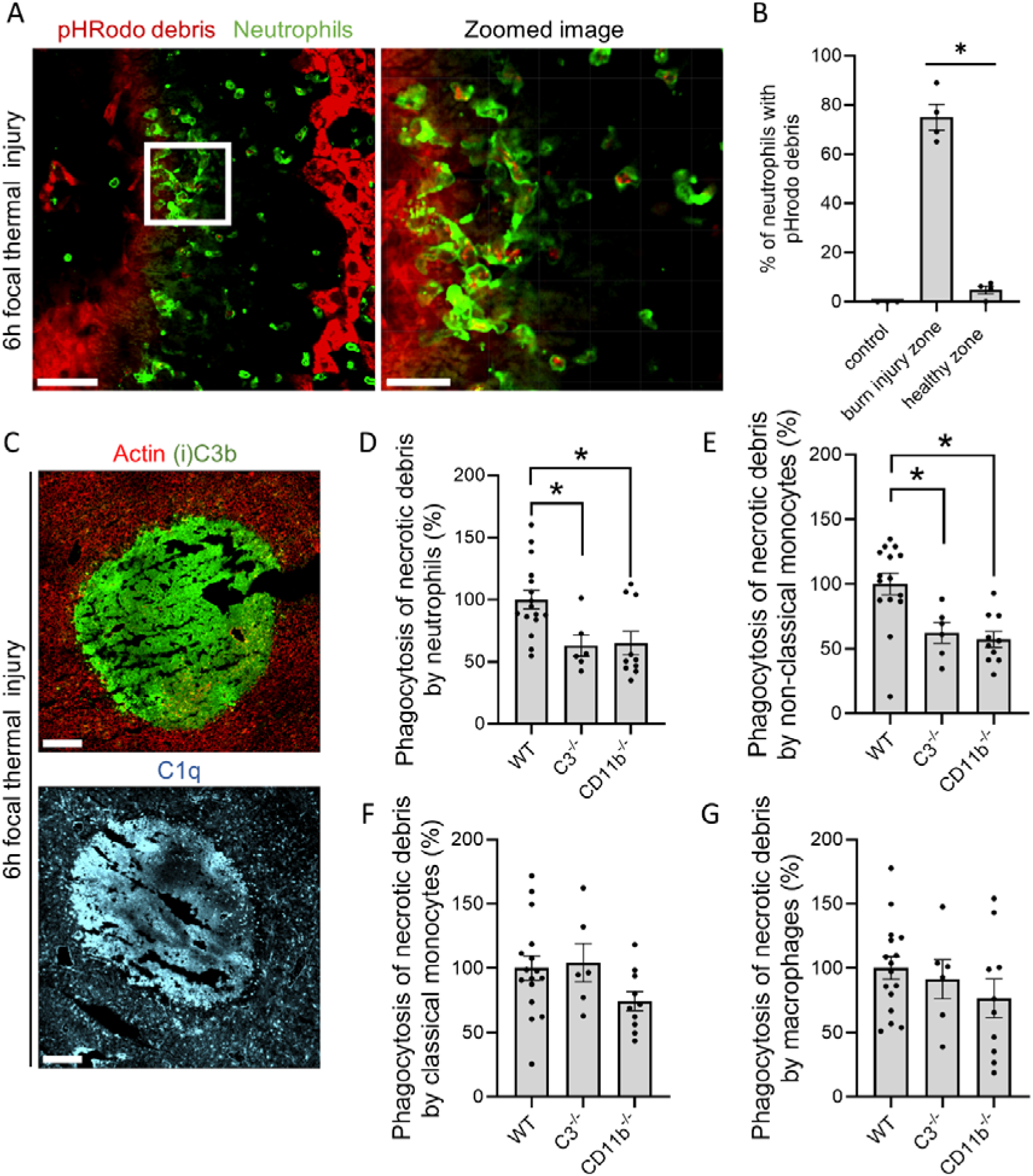
Complement opsonization and clearance of necrotic cell debris are not exclusive to drug-induced liver injury. (A) Representative image of neutrophils (green) and necrotic debris (pHRodo; red) in the liver 6h after FTI. pHRodo-SE was applied on top of the lesion 6h before imaging. Scale bar represents 50 µm, scale bar zoomed image represents 20 µm. (B) Quantification of the percentage neutrophils carrying pHRodo labeled debris at different areas of the injury. (C) Representative immunofluorescence stainings of liver cryosections 6h post burn injury. Green: (i)-C3b; Cyan: C1q. Scale bar represents 200 µm. (D-G) Flow cytometry of non-parenchymal cells isolated from liver FTI sites showing the percentage (D) pHrodo neutrophils (Ly6G), (E) pHrodo non-classical monocytes (Ly6C / CX_3_CR1), (F) pHrodo classical monocytes (Ly6C / CX_3_CR1 / CCR2) and macrophages (F4/80). Data are represented as mean ± SEM. Each dot represents a single mouse. *p≤0.05 compared to WT.

### Opsonization controls the volume of debris that is phagocytosed and regulates gene expression in neutrophils

Investigating the mechanisms driving debris phagocytosis *in vivo* is challenging due to the necessity of multiple knock-out strains and the technical limitations associated with observing cellular events in living mice. To overcome this, we developed an *in vitro* phagocytosis assay, enabling us to study the role of specific complement proteins in necrotic debris clearance. This approach also allowed us to verify our findings in human neutrophils and human necrotic debris. In this assay, necrosis was induced by mechanically disrupting HepG2 cells, whereafter sera lacking specific complement components were used to opsonize the debris. Images were taken with a confocal microscope 3h after combining the opsonized necrotic debris with human or mouse neutrophils. Importantly, the HepG2 debris itself does not contain C3/(i)C3b, as demonstrated by an *in vitro* HepG2 debris spot that failed to be immunostained by an antibody against C3/(i)C3b. However, the debris became clearly stained when opsonized with whole serum (**supplemental Figure 6A-B**), also confirming the capacity of debris to induce complement activation.

Engulfment of pHRodo-labeled necrotic debris (red) by live neutrophils can be visualized using 3D reconstruction of images from the phagocytosis experiment (**Figure 6A**). Interestingly, the percentage of bone-marrow derived mouse neutrophils that performed phagocytosis did not differ when presented with debris opsonized with serum, serum lacking C3 (from C3^-/-^ mice) or lacking antibodies (from Rag2^-/-^ mice) (**Figure 6B**). However, the volume of the necrotic debris internalized by neutrophils was significantly reduced when the debris was opsonized with serum lacking C3 or antibodies (**Figure 6C**). Likewise, freshly isolated neutrophils from healthy donors showed no difference in phagocytosis rates when debris was opsonized with serum, C1q-depleted serum or C3-depleted serum (**Figure 6D**). Nevertheless, a significant decrease in the volume of debris uptake was observed when it was opsonized with serum lacking C3 (**Figure 6E**). Latrunculin served as a negative control, as it inhibits phagocytosis globally by disrupting actin polymerization. These findings underscore the ability of both human and mouse neutrophils to internalize necrotic debris, while highlighting the effect of complement on the amount of debris taken up through phagocytosis.

**Figure 6.**
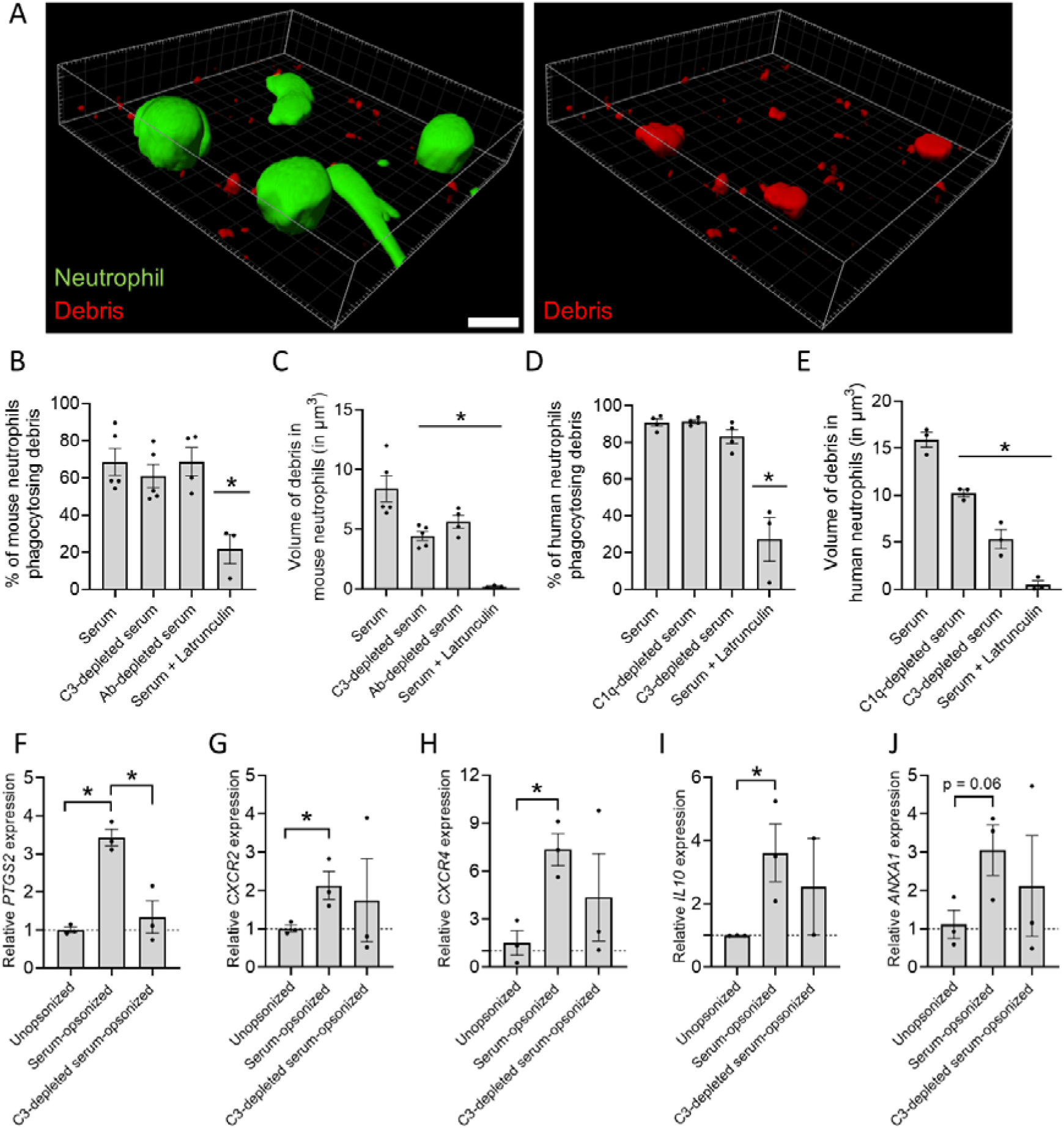
Opsonization controls the volume of debris phagocytosed and regulates gene expression in neutrophils. (A) 3D reconstruction image of human neutrophils stained with Calcein (green) phagocytosing pHrodo-labeled necrotic debris (red). Scale bar represents 10 µm. (B) Percentage of bone-marrow derived mouse neutrophils phagocytosing necrotic debris opsonized with serum, C3-depleted serum (from C3^-/-^ mice) or antibody-depleted serum (from Rag2^-/-^ mice). (C) Volume of necrotic debris phagocytosed by bone-marrow derived mouse neutrophils which was opsonized with serum, C3 serum or C1q serum. (D) Percentage of human neutrophils phagocytosing necrotic debris opsonized with serum, C1q-depleted serum or C3-depleted serum. (E) Volume of necrotic debris phagocytosed by human neutrophils which was opsonized with serum, C1q-depleted serum or C3-depleted serum. (F-J) Gene expression of human neutrophils incubated with unopsonized, serum-opsonized or C3-depleted serum-opsonized human necrotic debris. Data are normalized to the average expression of 3 housekeeping genes (GAPDH, 18s and CDKN1A) and represented as 2^−ΔΔCt^ relative to the unopsonized group. 10 µM latrunculin is used as a negative control. Data are represented as mean ± SEM. Each dot represents an independent experiment. *p≤0.05 compared to serum group or between indicated groups.

To investigate the impact of debris phagocytosis on gene expression, qPCR was performed on human neutrophils incubated with necrotic debris from HepG2 cells. Neutrophils were exposed to pure unopsonized debris, debris opsonized with normal serum or C3-depleted serum-opsonized debris for 3 hours. The data were normalized to unopsonized debris in order to account for stimulation by DAMPs present in necrotic cell debris. Interestingly, uptake of serum-opsonized debris induced the upregulation of *PTGS2* (encoding COX2) in neutrophils **(Figure 6F)**. COX2 is an enzyme with dual role in inflammation, catalyzing the production of pro-inflammatory prostaglandins from arachidonic acid, such as PGE2, but also participating in the synthesis of numerous pro-resolving lipid mediators.^32^ *PTGS2* upregulation was reversed when the debris was opsonized with serum lacking C3, indicating a direct effect of complement on the upregulation of COX2, which was already observed for monocytes but not neutrophils.^33^ *CXCR2*, coding for a major chemokine receptor in neutrophils that promotes both chemotaxis and reverse migration^34,35^ was also upregulated by incubation with opsonized debris, however, no difference in comparison with C3-depleted serum was observed **(Figure 6G)**. In addition, incubation with opsonized debris led to expression of other immunoregulatory and pro-resolving genes in neutrophils, namely, *CXCR4*, encoding a chemokine receptor associated with homing of neutrophils to the bone marrow for apoptosis and removal^36,37^; the immunoregulatory cytokine *IL10*, and *ANXA1*, encoding the protein annexin A1 that dampens leukocyte chemotaxis, respiratory burst and phagocytosis **(Figure 6H-J)** ^38^. The absence of C3 in the serum, however, did not impact the expression of *CXCR2*, *CXCR4*, *IL10 or ANXA1* by neutrophils. Moreover, alterations in gene expression in neutrophils are specific, since multiple genes were unaffected by stimulation with opsonized debris, including *ALOX5, CYBB, CASP3, ARG1* and *ABCA1* (**supplemental Figure 6C-G**). Overall, the gene expression induced by clearance of opsonized necrotic debris represents a pro-resolving response of neutrophils to debris, which is central in promoting tissue repair.

## Discussion

The composition of necrotic cell debris varies depending on the type of cell death as well as on the type of cell dying.^39^ To efficiently clear this diverse and complex intracellular material, we hypothesized that C3 could opsonize the debris and cause clearance by complement-mediated phagocytosis. Overall, the opsonophagocytic function of C3 has been demonstrated on apoptotic cells^40^, bacteria^5^, fungi^8^ and amyloid plaques^41^, as evidenced by a reduction in phagocytosis of these targets in C3^-/-^ mice. Unfortunately, limited research has explored its role in clearing necrotic debris from tissues, despite the value of preventing excessive inflammatory responses that delay recovery. *In vitro* studies have identified C1q and C3 as indispensable components in the phagocytosis of late apoptotic/necrotic cells by monocyte-derived macrophages.^22^ However, conclusive evidence in an *in vivo* context was lacking, primarily due to technical challenges of directly visualizing debris clearance *in vivo*, which we overcame by using appropriate mouse models and intravital microscopy with pH-dependent fluorescent dyes.

We observed specific deposition of C1q and C3b at necrotic lesions in the liver, in line with previous literature **(Figure 1)**.^43–45^ This phenomenon is not exclusive to the liver, as it has also been demonstrated in the brain in Alzheimer’s disease and multiple sclerosis^46,47^, heart attack^48^ and wounds (trauma, burns)^49^. Whether this deposition is beneficial or harmful to the course of injury is injury-dependent. When controlled, complement promotes tissue repair, while excessive activation causes uncontrolled inflammation and tissue damage.^50^ Further research is needed to clarify the precise role of complement in the context of specific injuries.

We demonstrated that in the absence of antibodies IgM and IgG, C1q still bound necrotic debris **(Figure 2)**, likely due to its interaction with various ligands which include histones, DNA, C-reactive protein, pentraxin 3 and serum amyloid P component.^4,51–53^ Interestingly, the absence of antibodies significantly reduced C3b deposition, even though Rag2^-/-^ mice have elevated C3 levels compared to WT mice.^29^ This suggests that C3b deposition on necrotic lesions relies on the classical complement pathway, which is hampered in the absence of antibodies despite C1q presence. An *in vitro* study corroborated this, since adding C1q to sera lacking IgM and C1q did not affect C3 deposition on apoptotic cells.^54^ Of course, the activation of the alternative and lectin complement pathway in response to necrotic cells should not be overlooked, as properdin, a positive regulator of the complement system, has been proven to bind to necrotic cells and activate the alternative pathway.^55^ Realization that the classical pathway is activated reveals additional opportunities to ameliorate debris clearance. Patients with severe necrotic injuries might benefit from intravenous immunoglobulin (IVIG) supplementation and blood transfusions. The administered natural antibodies may bind necrotic debris, triggering C3 cleavage via the classical pathway, and aiding in the clearance of debris. Moreover, our previous work showed that the supplementation of natural antibodies directly enhanced Fc receptor-mediated phagocytosis, meaning that debris would be cleared via both complement- and Ab-mediated phagocytosis.^25^

Evaluating liver injury in C3^-/-^ mice following APAP overdose showed us a delayed recovery and the prolonged accumulation of necrotic debris **(Figure 3)**. Roth *et al.*^43^ observed lower serum ALT levels in C3^-/-^ mice 6 and 12h post-APAP, possibly due to the administration of a lower dose of 300 mg/kg APAP intraperitoneally and the shorter evaluation time. However, other studies have similarly noted impaired liver regeneration in C3^-/-^ mice after toxic-injury induced by CCl and partial hepatectomy.^56,57^ Although impaired liver resolution and debris accumulation in C3^-/-^ mice could be explained by impaired debris phagocytosis, other factors also contribute to this worsened phenotype. The absence of anaphylatoxins C3a and C5a impact liver regeneration by affecting hepatocyte priming and inhibiting neutrophil and monocyte chemotaxis.^58,54^ In addition to the decreased presence of monocytes at the injury site to perform debris phagocytosis, the reduced differentiation into monocyte-derived macrophages also contributes to delayed injury resolution, as observed in CCR2^-/-^ mice.^59,60^ This emphasizes that liver resolution is a complex process involving multiple factors, with debris phagocytosis being just one of the contributing events.

APAP-induced liver injury is characterized by the accumulation of pro-inflammatory mediators such as TNF-α, CXCL1, CXCL2 and CXCL6, which attracts neutrophils.^61^ We showed that neutrophils, along with macrophages, rely on complement to phagocytose debris **(Figure 4I-J)**. This phenomenon is not limited to the liver, as similar uptake of debris occurred in the lungs of mice with acid-induced lung injury.^62^ Using confocal microscopy, we observed an average of two DNA-containing phagosomes per cell, consistent with previous findings of macrophages ingesting one or more small cytosolic particles from necrotic cells.^63^ Interestingly, cells that relied on complement for recruitment did not rely on it for phagocytosis, and *vice versa*, highlighting the existence of multiple pathways for leukocyte recruitment and debris clearance which may compensate each other.

To validate complement-dependent debris clearance, we used a liver FTI model that causes immediate cell death, preventing hepatocytes from undergoing programmed necrosis. In contrast, APAP overdose induces cell death at a slower rate, allowing mechanisms such as ferroptosis and necroptosis to be initiated.^30,31^ Although cell death in both models results in plasma membrane rupture and intracellular content release, the distinct molecular events preceding membrane damage can alter the nature of the necrotic remnants and, thus, the phagocytic signal that prompts their clearance. This means that observations from both models could differ due to the debris that is investigated (DNA versus protein), or due to the type of cell death that occurs (regulated versus unregulated). We observed the phagocytosis of pHRodo-labeled protein-rich necrotic debris, complementing findings of Wang *et al.* where neutrophils engulfed nuclear debris.^64^ Our results showed that phagocytosis in neutrophils and non-classical monocytes is complement-dependent in FTI **(Figure 5)**, adding a mechanistic layer to the role of CX_3_CR1^+^ monocytes in sterile injury resolution. Classical monocytes (CCR2^high^, CX_3_CR1^low^) surround the FTI site and transition into non-classical monocytes (CX_3_CR1^+^, CCR2^low^) essential for injury repair, a process dependent on IL-10 and IL-4.^65^

Human neutrophils exhibited a pro-resolving phenotype after phagocytosis of necrotic debris, marked by the gene upregulation of *IL10*, *PTGS2, CXCR4, CXCR2* and *ANXA1* **(Figure 6F-J)**. This gene expression shifts depend on the type of meal ingested, as illustrated in macrophages, where efferocytosis of apoptotic cells triggers an anti-inflammatory response.^66^ Research on efferocytosis revealed that the uptake of lipids from apoptotic bodies stimulates sterol receptors (PPARs and Liver X receptors), triggering an anti-inflammatory response via IL-10 and TGF-β production.^67,68^ Due to the lipid-rich nature of the debris, these insights could apply to necrotic cells as well. The upregulated expression of CXCR4 is associated with reverse migration of the neutrophils to the bone marrow, as similarly shown *in vivo* by Wang *et al*.^64^ This process would in theory assist to alleviate the burden of dead cells to be cleared at the injury site when neutrophils undergo apoptosis. The opsonization of debris in the absence of C3 had no impact on gene expression, with the exception of *COX2*. This suggests that phagocytosis itself and the content that is phagocytosed, rather than the specific opsonization process, is primarily responsible for driving the expression of genes associated to resolution.

The in vitro phagocytosis assay showed that both mouse and human neutrophils ingested necrotic debris, however, the phagocytosed volume was reduced in the absence of C1q and C3 **(Figure 6)**. Opsonins affect hydrophobicity and surface charge, consequently influencing receptor interactions.^69^ This could potentially impact debris removal and explain the larger area of necrotic debris observed in C3^-/-^ mice 48h post-APAP (**Figure 3D-E)**. Also, mechanosensing of the target, which drives actin-based protrusions to mediate particle internalization, might be impaired, possibly due to the reduced stiffness/rigidity of the debris in the absence of C3.^70^

In conclusion, our study demonstrates the crucial role complement proteins play in the opsonization and subsequent phagocytosis of necrotic debris. This mechanism was confirmed in both mouse and human neutrophils, irrespective of the nature of injury (chemical or thermal). This highlights a general complement-dependent pathway for debris clearance specifically for neutrophils. Consequently, individuals with complement deficiencies, whether it is due to genetic factors or auto-immune diseases, might exhibit impaired clearance of necrotic debris, causing prolonged inflammation and poor tissue regeneration. These individuals could potentially benefit from C3 or plasma supplementation to enhance debris clearance and fasten injury recovery.

## Acknowledgments

SV and SS hold PhD fellowships from the Research Foundation of Flanders (FWO-Vlaanderen; SB1S56521N and 1116922N, respectively). This work is supported by FWO-Vlaanderen Junior Research Grants (G058421N and G025923N), a KU Leuven C1 grant (14/23/143), a Global PhD Fellowship between KU Leuven and University of Maastricht (GPMU/22/006) and the Rega Foundation.

## Author contributions

SV and PEM designed the experiments. SV and PEM wrote the manuscript; SV, MSM, EB and SS conducted the experiments; SV, MSM, PP and PEM analyzed and discussed data.

## SUPPLEMENTAL FIGURE LEGENDS

**Supplemental Figure 1. Liver injury in response to paracetamol (APAP) overdose in mice.** (A) Serum ALT levels in mice 12, 24 and 48h after receiving an overdose of 600 mg/kg APAP. (B) Area of necrosis in the liver of mice 12, 24 and 48h after receiving an overdose of 600 mg/kg APAP, determined by histopathology. (C) Representative H&E staining images of a control liver and necrotic liver 12h after APAP administration. Scale bar represents 100 µm. Image quantifications were pooled from 10 fields of view. Data are represented as mean ± SEM. Each dot represents a single mouse. *p≤0.05 compared to control; #p≤0.05 between indicated groups. APAP = acetaminophen, ALT = alanine aminotransferase.

**Supplemental Figure 2. Time-response evaluation of C1q and C3b deposition at the sites of necrotic injury in the liver.** (A,B) Representative immunofluorescence images of liver cryosections from control mice and mice 24, 48 and 72 hours after receiving an overdose of acetaminophen (APAP; 600 mg/kg). Green: (i)C3b, red: fibrin(ogen), Orange: IgM; Cyan: C1q. Scale bar represents 50 µm.

**Supplemental figure 3. DNAse treatment validates the internalization of DNA debris in CD11b+ leukocytes.** (A-F) Flow cytometry of liver non-parenchymal cells identifying the percentage of Neutrophils (Ly6G^+^), monocytes (Ly6G^-^ / Ly6C^+^ / CCR2^+^) and macrophages (F4/80^+^) and the percentage of cells expressing CD11b in the injured liver 24h after APAP overdose. (G) Representative IVM images of the injured liver 24h after an APAP overdose. 2 µl of the cell-impermeable DNA dye Sytox Green was injected 1h before imaging. Leukocytes were labeled with anti-CD11b antibody (Red). 1 mg DNAse I was injected intravenously to remove extracellular DNA. Scale bar represents 50 µm, scale bar zoomed image represents 10 µm.

**Supplemental figure 4. Quantification and visualization of leukocytes and complement proteins in the focal thermal injury of the liver.** (A) Representative image of a 6h burn injury showing the different injury zones (core of injury, burned and healthy areas). Green: neutrophils; Red, pHRodo. Scale bar represents 200 µm. (B-C) Mean fluorescence intensity of pHRodo-labeled necrotic debris in neutrophils and classical monocytes in healthy and burned areas. (D) Representative immunofluorescence images of liver cryosections from 6h burn injury. Green: (i)-C3b; Cyan: C1q. Scale bar represents 150 µm. (E) Representative immunofluorescence images of liver cryosections from 6h burn injury. Green: (i)-C3b; Cyan: Fibrin(ogen); Red: f-actin. Scale bar represents 150 µm. (F-I) Flow cytometry of liver non-parenchymal cells identifying neutrophils (Ly6G^+^), classical monocytes (Ly6C^+^ / CX_3_CR1^-^ / CCR2^+^), non-classical monocytes (Ly6C^+^ / CX_3_CR1^-^ / CCR2^+^) and macrophages (F4/80^+^) 6h after burn in WT, C3^-/-^ and CD11b^-/-^ livers. Data are represented as mean ± SEM. Each dot represents a mouse. *p≤0.05.

**Supplemental figure 5. CD11b does not participate in the recovery from drug-induced liver injury** (A) ALT levels of WT and CD11b^-/-^ mice challenged with APAP for 24 or 48h. (B) Quantification of the fibrin(ogen)^+^ area fraction in liver cryosections of WT and CD11b^-/-^ mice challenged with APAP in experiment (A). (C) ALT levels of mice challenged with APAP, treated with 40 µg isotype control or 40 µg anti-CD11b blocking antibody 6 and 12h post-APAP. Samples were collected at either 24 or 48h post APAP overdose. (D) Quantification of the fibrin(ogen)^+^ area fraction in liver cryosections of experiment (C). (E) Percentage of neutrophils expressing CD11b determined by flow cytometry of experiment (C).

**Supplemental figure 6. Complement activation on purified necrotic debris in vitro** (A) Representative images of necrotic debris form HepG2 cells opsonized with 20% serum or heat-inactivated serum. DNA debris is labeled with Hoechst (blue) and C3/C3b/iC3b in Cyan. Scale bar represent 100 µm. (B) Mean fluorescence intensity of C3/C3b/iC3b labeling on in vitro necrotic debris opsonized with 20% serum or heat-inactivated (HI) serum. (C-G) Gene expression of human neutrophils incubated with unopsonized, serum-opsonized or C3-depleted serum-opsonized human necrotic debris. Data is normalized to the average expression of 3 housekeeping genes (GAPDH, 18s and CDKN1A) and represented as 2^−ΔΔCt^ relative to the unopsonized group. Images were taken with a Zeiss Axiovert 200M fluorescence microscope and analyzed with FIJI. Data are represented as mean ± SEM. *p≤0.05.

