## Supplemental data for "Complement activation at injury sites drives the phagocytosis of necrotic cell debris and resolution of liver injury"

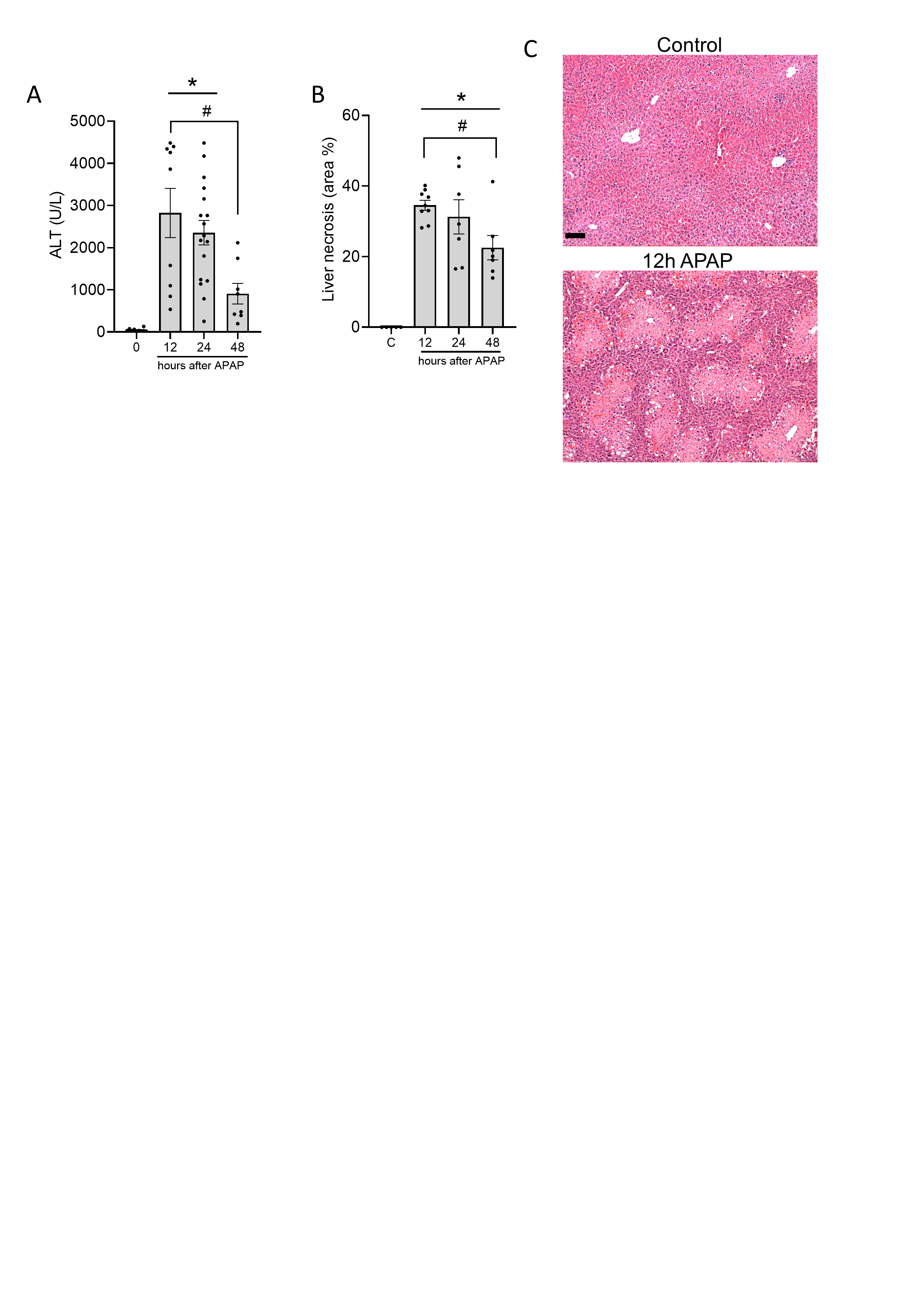

**Supplemental Figure 1. Liver injury in response to paracetamol (APAP) overdose in mice.** (A) Serum ALT levels in mice 12, 24 and 48h after receiving an overdose of 600 mg/kg APAP. (B) Area of necrosis in the liver of mice 12, 24 and 48h after receiving an overdose of 600 mg/kg APAP, determined by histopathology. (C) Representative H&E staining images of a control liver and necrotic liver 12h after APAP administration. Scale bar represents 100 µm. Image quantifications were pooled from 10 fields of view. Data are represented as mean ± SEM. Each dot represents a single mouse. *p≤0.05 compared to control; #p≤0.05 between indicated groups. APAP = acetaminophen, ALT = alanine aminotransferase.

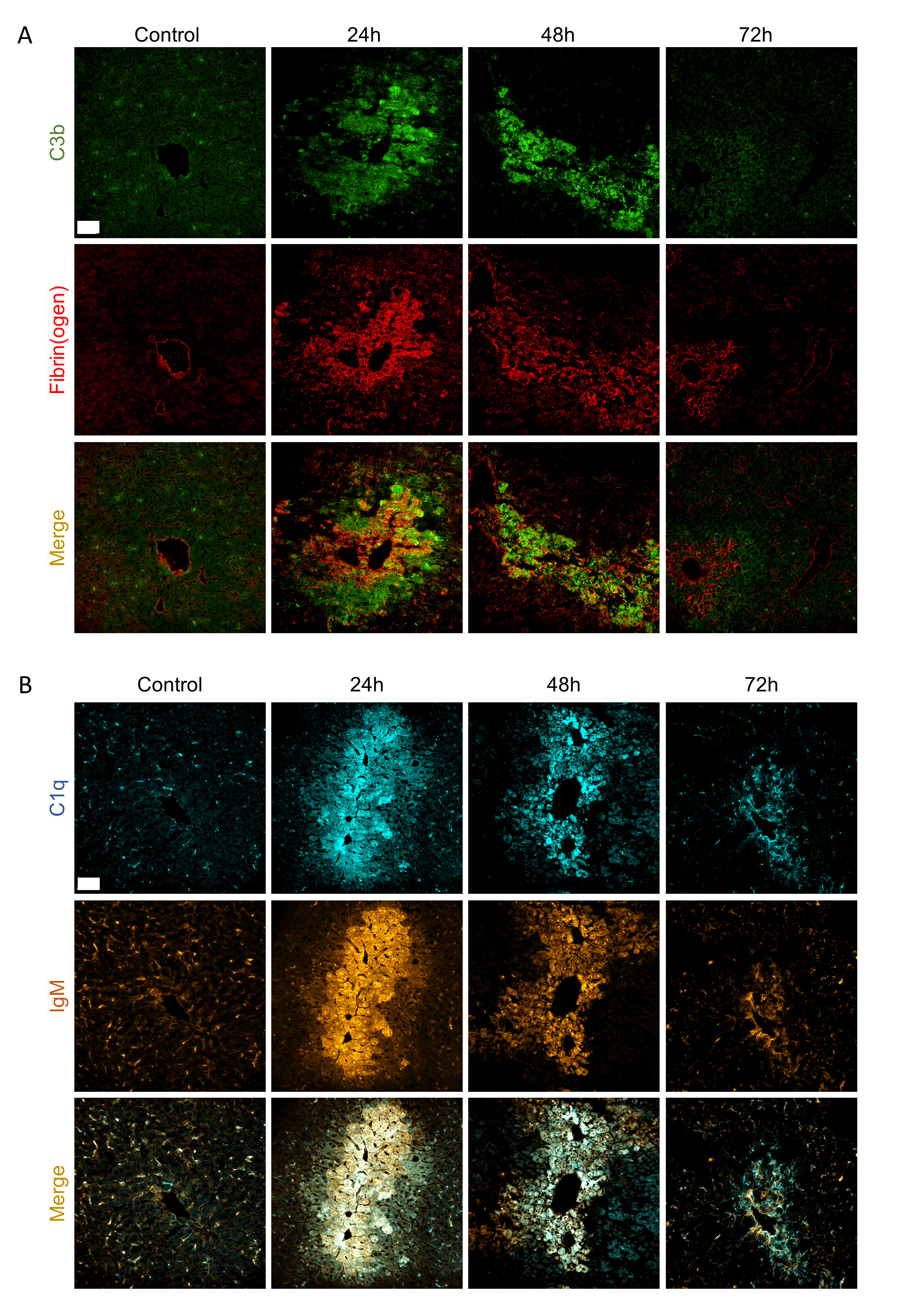

**Supplemental Figure 2. Time-response evaluation of C1q and C3b deposition at the sites of necrotic injury in the liver.** (A,B) Representative immunofluorescence images of liver cryosections from control mice and mice 24, 48 and 72 hours after receiving an overdose of acetaminophen (APAP; 600 mg/kg). Green: (i)C3b, red: fibrin(ogen), Orange: IgM; Cyan: C1q. Scale bar represents 50 µm.

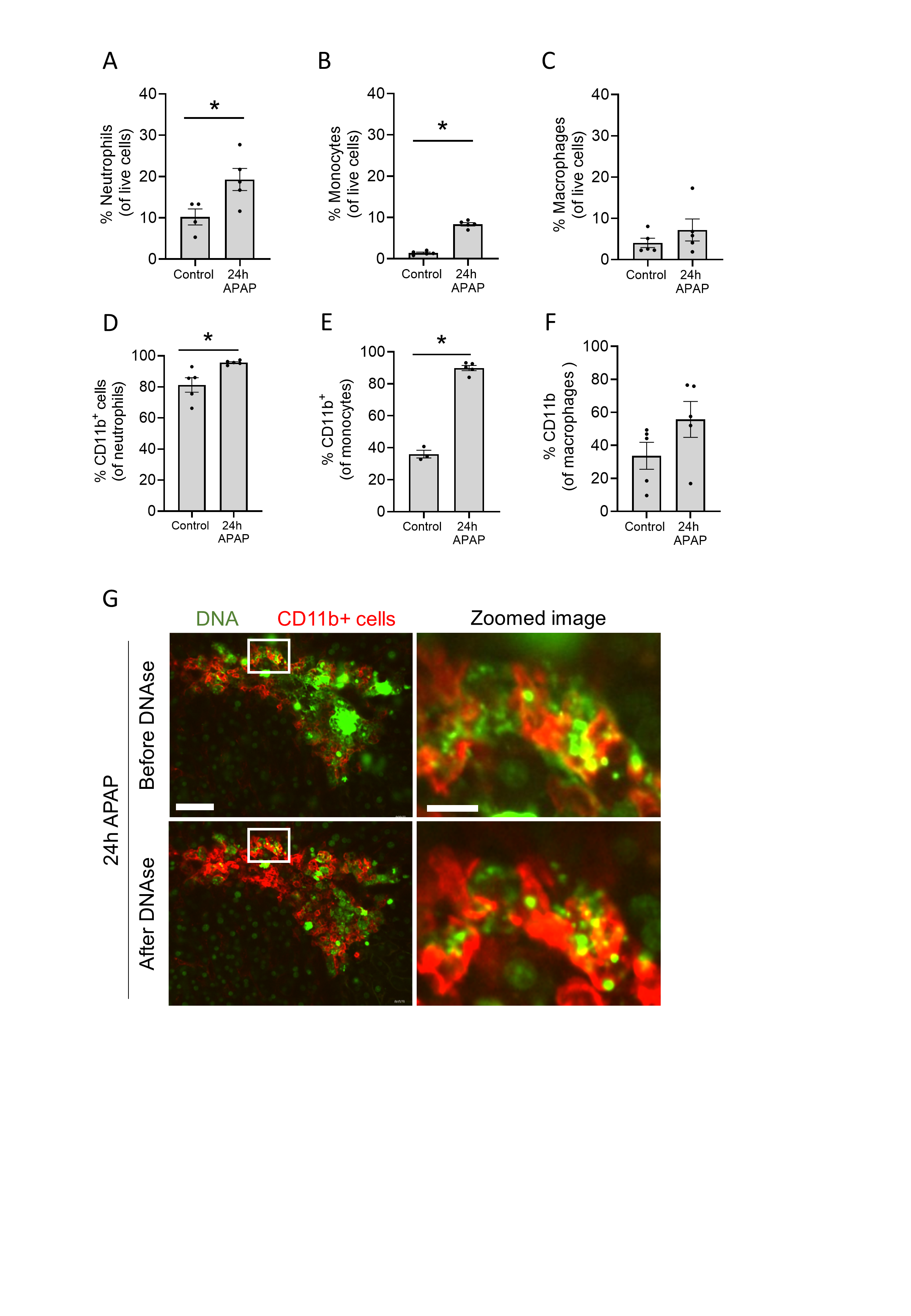

**Supplemental figure 3. DNAse treatment validates the internalization of DNA debris in CD11b+ leukocytes.** (A-F) Flow cytometry of liver non-parenchymal cells identifying the percentage of Neutrophils (Ly6G^+^), monocytes (Ly6G^-^ / Ly6C^+^ / CCR2^+^) and macrophages (F4/80^+^) and the percentage of cells expressing CD11b in the injured liver 24h after APAP overdose. (G) Representative IVM images of the injured liver 24h after an APAP overdose. 2 µl of the cell-impermeable DNA dye Sytox Green was injected 1h before imaging. Leukocytes were labeled with anti-CD11b antibody (Red). 1 mg DNAse I was injected intravenously to remove extracellular DNA. Scale bar represents 50 µm, scale bar zoomed image represents 10 µm.

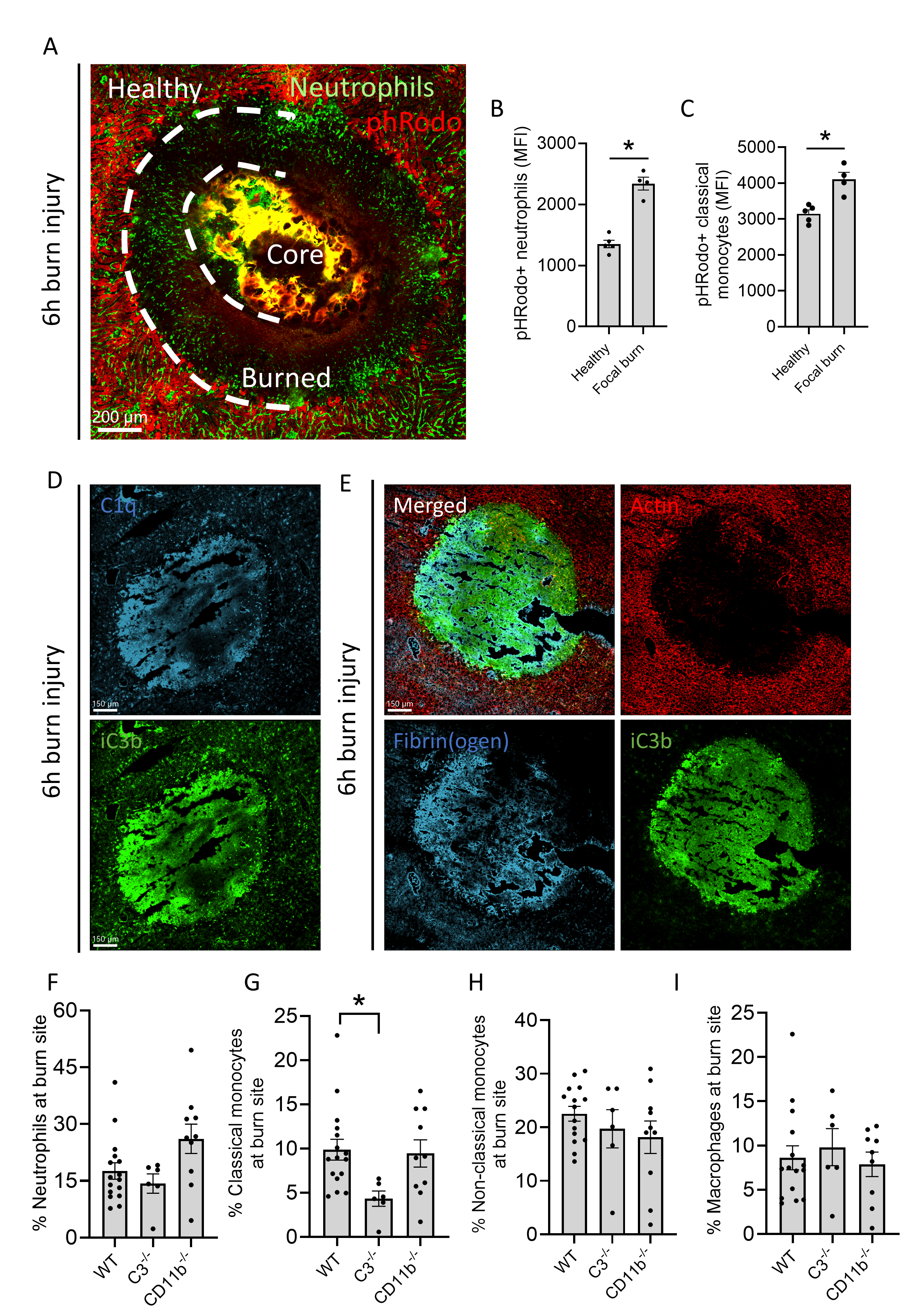

**Supplemental figure 4. Quantification and visualization of leukocytes and complement proteins in the focal thermal injury of the liver.** (A) Representative image of a 6h burn injury showing the different injury zones (core of injury, burned and healthy areas). Green: neutrophils; Red, pHRodo. Scale bar represents 200 µm. (B-C) Mean fluorescence intensity of pHRodo-labeled necrotic debris in neutrophils and classical monocytes in healthy and burned areas. (D) Representative immunofluorescence images of liver cryosections from 6h burn injury. Green: (i)-C3b; Cyan: C1q. Scale bar represents 150 µm.
(E) Representative immunofluorescence images of liver cryosections from 6h burn injury. Green: (i)-C3b; Cyan: Fibrin(ogen); Red: f-actin. Scale bar represents 150 µm. (F-I) Flow cytometry of liver non-parenchymal cells identifying neutrophils (Ly6G^+^), classical monocytes (Ly6C^+^ / CX_3_CR1^-^ / CCR2^+^), non-classical monocytes (Ly6C^+^ / CX_3_CR1^-^ / CCR2^+^) and macrophages (F4/80^+^) 6h after burn in WT, C3^-/-^ and CD11b^-/-^ livers. Data are represented as mean ± SEM. Each dot represents a mouse. *p≤0.05.

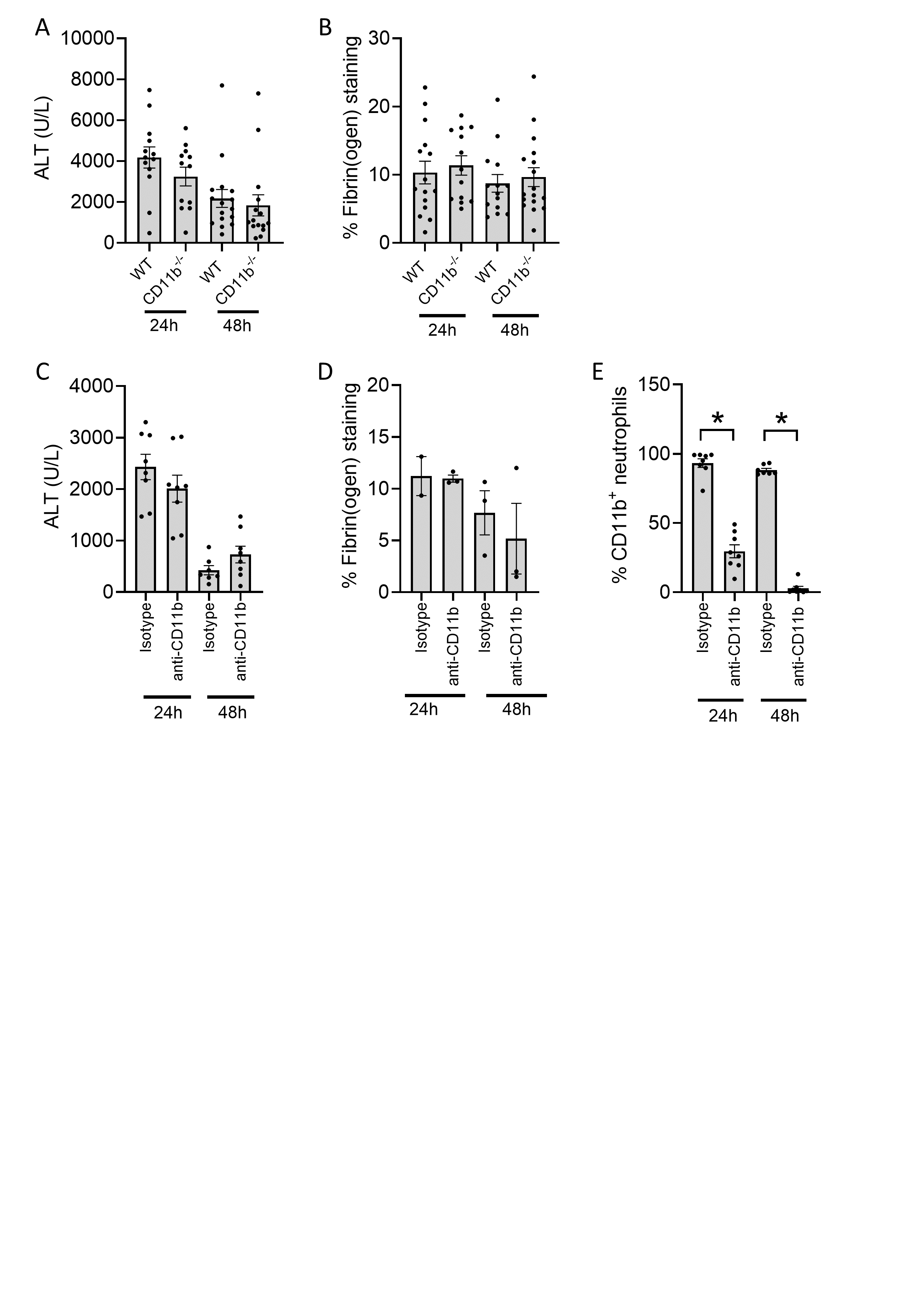

**Supplemental figure 5. CD11b does not participate in the recovery from drug-induced liver injury** (A) ALT levels of WT and CD11b^-/-^ mice challenged with APAP for 24 or 48h. (B) Quantification of the fibrin(ogen)^+^ area fraction in liver cryosections of WT and CD11b^-/-^ mice challenged with APAP in experiment (A). (C) ALT levels of mice challenged with APAP, treated with 40 µg isotype control or 40 µg anti-CD11b blocking antibody 6 and 12h post-APAP. Samples were collected at either 24 or 48h post APAP overdose. (D) Quantification of the fibrin(ogen)^+^ area fraction in liver cryosections of experiment (C). (E) Percentage of neutrophils expressing CD11b determined by flow cytometry of experiment (C).

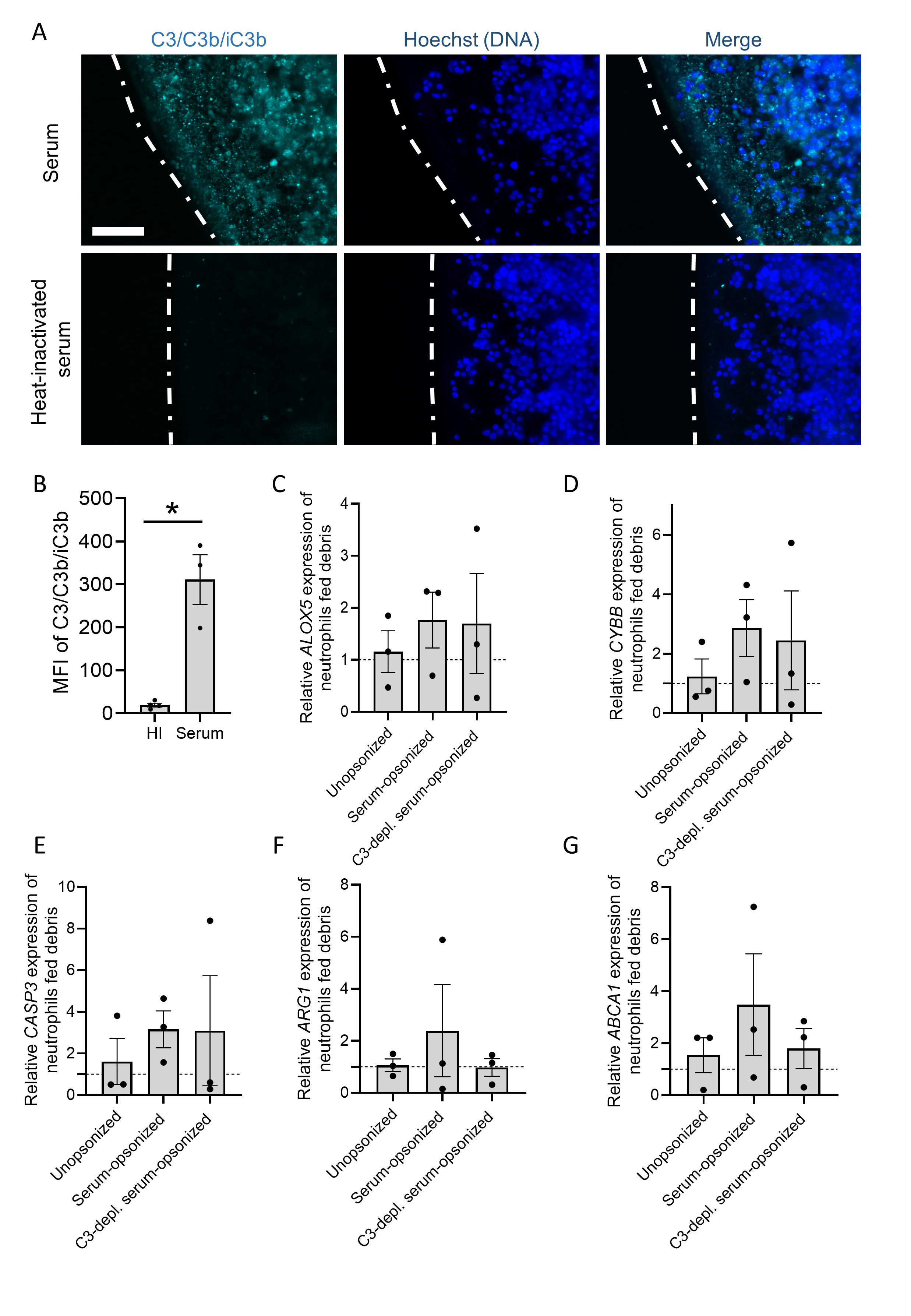

**Supplemental figure 6. Complement activation on purified necrotic debris *in vitro***
(A) Representative images of necrotic debris form HepG2 cells opsonized with 20% serum or heat-inactivated serum. DNA debris is labeled with Hoechst (blue) and C3/C3b/iC3b in Cyan. Scale bar represent 100 µm. (B) Mean fluorescence intensity of C3/C3b/iC3b labeling on in vitro necrotic debris opsonized with 20% serum or heat-inactivated (HI) serum. (C-G) Gene expression of human neutrophils incubated with unopsonized, serum-opsonized or C3-depleted serum-opsonized human necrotic debris. Data is normalized to the average expression of 3 housekeeping genes (GAPDH, 18s and CDKN1A) and represented as 2^–∆∆Ct^ relative to the unopsonized group. Images were taken with a Zeiss Axiovert 200M fluorescence microscope and analyzed with FIJI. Data are represented as mean ± SEM. *p≤0.05.
